# Tailored perception: listeners’ strategies for perceiving speech fit their individual perceptual abilities

**DOI:** 10.1101/263079

**Authors:** Kyle Jasmin, Fred Dick, Lori Holt, Adam Tierney

**Affiliations:** Department of Psychological Sciences, Birkbeck, University of London, London, UK; Institue of Cognitive Neuroscience, UCL, London, UK; Department of Experiment Psychology, UCL, London, UK; Department of Psychology, Carnegie Mellon University, Pennsylvania, USA

**Keywords:** Speech, Music, Amusia, Pitch, duration, prosody

## Abstract

In speech, linguistic information is conveyed redundantly by many simultaneously present acoustic dimensions, such as fundamental frequency, duration and amplitude. Listeners show stable tendencies to prioritize these acoustic dimensions differently, relative to one another, which suggests individualized speech perception ‘strategies’. However, it is unclear what drives these strategies, and more importantly, what impact they have on diverse aspects of communication. Here we show that such individualized perceptual strategies can be related to individual differences in perceptual ability. In a cue weighting experiment, we first demonstrate that individuals with a severe pitch perception deficit (congenital amusics) categorize linguistic stimuli similarly to controls when their deficit is unrelated to the main distinguishing cue for that category (in this case, durational or temporal cues). In contrast, in a prosodic task where pitch-related cues are typically more informative, amusics place less importance on this pitch-related information when categorizing speech. Instead, they relied more on duration information. Crucially, these differences in perceptual weights were observed even when pitch-related differences were large enough to be perceptually distinct to amusic listeners. In a second set of experiments involving musical and prosodic phrase interpretation, we found that this reliance on duration information allowed amusics to overcome their perceptual deficits and perceive both speech and music successfully. These results suggest that successful speech - and potentially music - comprehension is achieved through multiple perceptual strategies whose underlying weights may in part reflect individuals’ perceptual abilities.

## Introduction

### Speech perception as categorization

Speech perception requires that listeners segment and classify what they hear, mapping the continuous acoustic variation produced by the vocal system onto linguistic categories. This process occurs at and among multiple putative levels simultaneously, as smaller units such as phonemes, syllables, and words are combined into larger structures such as phrases, clauses and sentences (Bentrovato, Devescovi, D’Amico, Wicha, & Bates, 2003; Elman, 2009; Lerdahl & Jackendoff, 1985; McMurray, Aslin, Tanenhaus, Spivey, & Subik, 2008; Utman, Blumstein, & Burton, 2000). The acoustic cues that signal a particular linguistic category can be quite subtle, and are often distorted or degraded by noisy environments or competing talkers, making auditory classification during speech perception a challenging perceptual feat (Holt & Lotto, 2008).

Given that phonetic and prosodic categorization require precise and rapid detection of acoustic cues, one might think that biologically-based difficulties with auditory perception would severely impair speech perception. Individual differences in auditory processing are indeed highly prevalent (Kidd, Watson, & Gygi, 2007): listeners differ widely on many dissociable auditory skills, such as their degree of temporal and spectral resolution. However, decades of research have consistently found that non-verbal auditory skills either do not correlate - or only weakly correlate - with speech perception ability in normal-hearing adults, and factor analysis consistently separates verbal and non-verbal sound perception into separate factors (Karlin, 1942; Kidd et al., 2007; Stankov & Horn, 1980; Surprenant & Watson, 2001; Watson & Kidd, 2002; Watson, Jensen, Foyle, Leek, & Goldgar, 1982).

How can a listener both be severely impaired in auditory perception and yet successful in speech perception? One possible explanation is that the neurocognitive mechanisms for speech perception and non-verbal auditory perception are dissociable. According to motor theories of speech perception (Liberman & Mattingly, 1985; Liberman, Cooper, Shankweiler, & Studdert-Kennedy, 1967), speech perception takes place via motor simulation, a mechanism unavailable to the listener when perceiving nonspeech sounds. An alternate explanation is that speech is robust to individual differences in auditory perception *because it is structurally redundant*: multiple acoustic dimensions convey information about a given category. This property may make speech robust not only to external background noise (Winter, 2014) but also to internal “noise” that is thought to relate to imprecise representation of an auditory dimension (Patel, 2014). This raises the possibility that listeners may be able to compensate for impairments along a single auditory dimension by assigning greater perceptual weight to auditory dimensions that they are better able to process.

### Structural redundancy in speech

As mentioned above, speech is a highly redundant signal wherein a linguistic category may be conveyed by many acoustic cues. The distinction between voiced and unvoiced consonants (for example ‘rapid’ versus ‘rabid’) is conveyed by as many as 16 different cues (Lisker, 2016), the most prominent of these being the voice onset time (VOT, e.g., the time lapsed prior to voicing onset) and fundamental frequency (F0) (Haggard, Ambler, & Callow, 1970; Lisker, 1957; Massaro & Cohen, 1976). Acoustic redundancy is not limited to phonetic perception; it is a widespread feature of speech that occurs across multiple time scales and applies to many different linguistic features. These include *syllable stress* (‘PREsent’ versus ‘preSENT’), which is conveyed by increases in syllabic amplitude, duration, and F0 (Fear, Cutler, & Butterfield, 1995; Mattys, 2000), *linguistic focus* (‘it was HER’ versus ‘it WAS her’) conveyed by increases in word duration and greater F0 variation (Breen, Fedorenko, Wagner, & Gibson, 2010; Chrabaszcz, Winn, Lin, & Idsardi, 2014), and *phrase structure* (‘No dogs are here’ versus ‘No, dogs are here’) conveyed by F0 variation and increased duration for the syllable just before the phrase boundary (de Pijper & Sanderman, 1994; Streeter, 1978).

How do listeners integrate redundant information across different acoustic dimensions? Prior research focusing on the “average listener” (as aggregated over the behavior of groups of participants) has indicated that the multiple dimensions contributing to speech categorization are not perceptually equivalent (Breen et al., 2010; Haggard et al., 1970; R. Liu & Holt, 2015; Massaro & Cohen, 1976; McMurray & Jongman, 2011). Some acoustic dimensions more reliably and robustly signal speech category membership, and therefore carry more perceptual weight (“primary” dimensions) whereas other dimensions are somewhat less diagnostic and carry less perceptual weight (“secondary” dimensions). For example, when making perceptual decisions about voiced and unvoiced consonant-vowels in clear speech, VOT of the initial consonant is primary, while F0 of the following vowel is secondary (Haggard et al., 1970; Massaro & Cohen, 1976; 1977). Conversely, for linguistic focus, F0 is primary while word duration is secondary (Breen et al., 2010; Chrabaszcz et al., 2014).

However, perceptual weights are not fixed: dimensional weights shift when the fidelity of the signal is degraded (for review see Holt, Tierney, Guerra, Laffere, & Dick, 2018). For example, when speech is presented in masking noise, VOT is harder to detect, making F0 more effective at signaling voicing categories. Correspondingly, when listeners are asked to categorize speech in noise, F0 is up-weighted and VOT is down-weighted (Holt et al., 2018; Winn, Chatterjee, & Idsardi, 2013). Furthermore, dimensional weights also change when the co-occurrence statistics of different dimensions are shifted, as in a foreign accent. When VOT and F0 cues are presented in a manner that goes against their usual covariation in English (Idemaru & Holt, 2011; R. Liu & Holt, 2015), listeners down-weight reliance on the secondary (F0) dimension (Idemaru & Holt, 2011; 2014). In addition, experimentally increasing the variability of a given acoustic dimension (thereby making it less reliable) leads to down-weighting of the dimension (Holt & Lotto, 2006) and categorization training with feedback also can lead to changes in dimensional weights (Francis, Baldwin, & Nusbaum, 2000; Francis, Kaganovich, & Driscoll-Huber, 2008; S.-J. Lim & Holt, 2011). Overall, these results suggest that listeners continuously monitor and assess the evolving relationships among acoustic dimensions and speech categories, and adjust perceptual weights of acoustic dimensions accordingly (Holt et al., 2018; Holt & Lotto, 2006; Toscano & McMurray, 2010).

### Individual differences in dimensional weighting in perceptual categorization

Prior research on speech categorization has focused largely on perceptual weighting strategies common across groups of listeners with similar histories of speech experience (see studies discussed above). However, large individual differences (Yu & Zellou, 2019) in perceptual weighting have been reported across a variety of phonetic contrasts, including Mandarin tone (Chandrasekaran, Sampath, & Wong, 2010), place of articulation (Hazan & Rosen, 1991), stop consonant length (Idemaru, Holt, & Seltman, 2012), vowels (Kim, Clayards, & Goad, 2018). Moreover, individual differences in dimensional weighting reflect stable listening strategies rather than random variation, having been found to be similar across testing sessions (Idemaru et al., 2012; Kim et al., 2018; Kong & Edwards, 2016). These and many other studies have established that different listeners use different strategies when perceiving speech, with some listeners heavily weighting sources of evidence that others rely upon little.

What is poorly understood is the underlying cause of these differences. Considered in the context of the large individual differences in auditory perception reviewed above, an ideal strategy for one listener may be sub-optimal for another due to individual differences in the reliability of auditory perception. We hypothesize that one important factor driving individual differences in perceptual weighting is variability in the relative utility of different perceptual dimensions across listeners. Applying this hypothesis to an extreme case, individuals with a severe perceptual deficit specific to a particular auditory dimension may learn, over time, that this dimension is not useful for them, and perceptually weight it less in categorization - *even in cases when changes in that dimension are easily perceptible*. They instead may strategically weight redundant cues as an alternative route to effective perceptual categorization. Here we tested this hypothesis by investigating a population with a severe domain-general perceptual deficit for pitch: participants with congenital amusia.

### Speech perception in amusia

Congenital amusia provides an ideal test case for examining the consequences of deficits in the perception of a single acoustic dimension on perceptual weighting strategies. Amusia is a disorder affecting around 1.5% of the population (Peretz & Vuvan, 2017), which is characterized by problems detecting small changes in pitch (Hyde & Peretz, 2004; Vuvan, Nunes-Silva, & Peretz, 2015) in the absence of impairments in general cognitive or other perceptual skills. This impaired pitch perception leads to deficits in musical abilities such as singing and melody recognition (Ayotte, Peretz, & Hyde, 2002; Dalla Bella, Giguère, & Peretz, 2009) as well as linguistic abilities such as statement/question judgments and perception of tone-language speech in background noise (Ayotte et al., 2002; Hutchins, Gosselin, & Peretz, 2010; Jiang, Hamm, Lim, Kirk, & Yang, 2010; Nan, Sun, & Peretz, 2010; Patel, Foxton, & Griffiths, 2005; Patel, Wong, Foxton, Lochy, & Peretz, 2008; Peretz et al., 2002; Vuvan et al., 2015). Asking amusics about their experiences with language, however, reveals a paradox: only 7% of amusics report problems with speech perception in everyday life (F. Liu, Patel, Fourcin, & Stewart, 2010). How is it that amusics show problems with speech perception in the laboratory but self-report being unaffected in ecologically valid settings? One possibility is that amusics compensate for their poor pitch fidelity by relying on other, redundant, acoustic cues in the speech signal.

Here, we used amusia as a test case to examine whether impairment along a single perceptual dimension leads to a shift in how different sources of information are weighted in perceptual categorization. In Experiment 1, participants with amusia and control participants categorized stimuli drawn from a two-dimensional acoustic space that varied in the extent to which fundamental frequency of the voice (referred to as F0, a strong indicator of vocal pitch), as well as durational information, pointed toward one of two linguistic interpretations. In a *phonetic* paradigm, stimuli differed in the identity of a word-initial consonant whereas in a *prosodic* paradigm, stimuli differed in the location of linguistic focus. Crucially, the “primary” acoustic dimension for the phonetic experiment was durational in nature, but the primary dimension for the focus component was F0 – the dimension that amusics hear with less fidelity. We hypothesized that if an individual’s perception of a primary dimension is unimpaired, they should have no need to shift reliance to other, redundant perceptual dimensions because the primary dimension provides a robust signal to category identity (Holt et al., 2018; Schertz, Cho, Lotto, & Warner, 2015a; Wu & Holt, 2018). However, if perception of a primary dimension is indeed impaired, listeners may strategically down-weight it in favor of a secondary dimension for which perception is not impaired.

The stimulus sets fully sampled an acoustic space created by sampling the orthogonal duration and F0 dimensions with equal probability. Participants therefore heard some stimuli that clearly belonged to one category or the other, and other stimuli that were perceptually ambiguous. This allowed us to measure the perceptual weight of the F0 and duration dimensions separately for the phonetic and prosodic components of the experiment (Holt & Lotto, 2006). Critically, we confirmed with psychophysical testing that F0 differences among our stimuli were large enough to be perceptible *even to amusic participants*. This ensured amusics could potentially use F0 information in the task.

## Experiment 1

Experiment 1 tested the hypothesis that the extent to which individuals rely on particular acoustic dimensions in perceiving speech can be predicted by the informativeness of a given dimension (i.e. which one is primary) and the presence or absence of a perceptual deficit for that dimension. Participants assigned tokens of spoken language to linguistic categories based on a phonetic contrast for which the primary cue was related to duration (for which no participant had a perceptual deficit). *Phonetic* perception was assessed using synthesized speech sounds differing in voicing (/b/ versus /p/), which can be conveyed via differences in voice onset time (VOT; a duration-related cue) and fundamental frequency of the following vowel (F0; a pitch-related cue). As described above, most listeners rely predominantly on VOT in /b/-/p/ categorization, and F0 is secondary (Haggard et al., 1970)). We predict that performance on this task should not differ between groups; none of the participants were impaired in perceiving the primary dimension, and could therefore rely upon it. The same participants also completed a similar task that examined a *prosodic* contrast for which the primary dimension was, instead, F0 (related to pitch, for which amusics have a deficit) and duration was secondary. Here, categorization of prosodic focus was primarily conveyed via pitch accents (carried by F0) and, to a lesser extent, durational lengthening (Breen et al., 2010). In this case, amusics are impaired at the primary, F0, dimension. Thus, we predict that they will perceptually weight F0 less than control participants *even across F0 differences than are easily perceptible for amusic participants,* based on their pitch thresholds. This would indicate that any group differences in perceptual cue weighting between amusics and controls reflect differences in categorization strategy rather than simply an inability to detect the cues.

### Methods

#### Participants

Sixteen amusics (10 F, age = 60.2 +- 9.4) and 15 controls (10 F, age = 61.3 +- 10.4) were recruited from the UK and were native British English speakers with the exception of one amusic whose native language was Finnish but acquired English at age 10. (This subject was excluded from the Linguistic Phrase and Focus Test analyses of Experiment 2). Experiment 1 participants (11 amusic (6 F, age = 59.3), 11 control (8 F, age = 60.4)) were a subset of the full sample. All participants gave informed consent and ethical approval was obtained from the ethics committee for the Department of Psychological Sciences, Birkbeck, University of London. Participants were compensated £10 per hour of participation. Amusia status was obtained using the Montreal Battery for the Evaluation of Amusia (MBEA), in which participants heard pairs of short melodic phrases and rhythms across 5 subtests -- Contour, Scale, Interval, Rhythm, Meter, and Musical Memory -- and judged whether each pair were identical or different. Participants with a composite score (summing the Scale, Contour and Interval tests scores) of 65 or less were classified as amusic (Peretz, Champod, & Hyde, 2003). No participant in either group had extensive musical experience.

#### Phonetic Cue Weighting

Each phonetic block consisted of repetitions of a single word, spoken by a female American English speaker, that varied from “beer” (IPA: /bier/) to “pier” (IPA: /pier/) along two orthogonal acoustic dimensions. Praat 5.0 (Boersma, 2002) was used to alter original recordings of the words “beer” and “pier” such that the voice onset time (VOT) ranged from - 5 ms to 15 ms in 5 ms increments. The F0 onset frequency of the vowel was manipulated manually in Praat such that it varied from 200 to 320 in 30 Hz increments. The F0 remained at this frequency for 80 ms, after which it decreased linearly to 180 Hz over the following 150 ms. Consistent with the distributional regularities of English, shorter VOT and lower initial F0 sounded more like /b/ while longer VOT and higher initial F0 sounded like /p/ (Idemaru & Holt, 2011). A full description of the stimulus creation methods can be found in Idemaru & Holt (2011). Each of the 5 F0 levels was crossed with each of the 5 VOT (duration) levels to make 25 stimuli. The stimulus set can be found in the online materials.

To ensure that the intervals along the F0 dimension were large enough for amusics to detect, we measured the median F0 of the vowel following the initial stop consonant for each of the five levels of F0 morphing. The vowel F0 for the five levels was 200, 230, 260, 290, and 320 Hz, with a mean difference between adjacent F0 levels of 2 semitones, well above the pitch discrimination threshold of all participants, which did not exceed 1.5 semitones (see Experiment 2).

#### Prosodic Cue Weighting

Linguistic focus materials were constructed from combinations of two phrases, “Dave likes to STUDY music” or “Dave likes to study MUSIC” (stress indicated in uppercase). These phrases were spoken by an actor who read two compound sentences with an intervening conjunction, e.g. “Dave likes to STUDY music, but he doesn’t like to PLAY music” and “Dave likes to study MUSIC, but he doesn’t like to study HISTORY”. The actor was asked to place contrastive accents to emphasize the capitalized words. These recordings were then cropped, leaving only the initial phrases listed above. Using STRAIGHT software (Kawahara & Irino, 2005), the two versions were manually time aligned. We then produced a set of 49 different stimuli by varying the extent to which F0 and durational information supported one versus the other focus interpretation. The duration and F0 information disambiguating the ‘focused’ word varied from 0% to 100% morphing rates for F0 and for duration, in 17% increments (0%, 17%, 33%, 50%, 67%, 83%, 100%). Because the dimensions were fully crossed, some combinations of F0 and duration cued an interpretation jointly, others conflicted, and tokens near the center of the space were more perceptually ambiguous (Fig. 1). Examples of the stimuli are provided in online materials.

**Figure 1:**
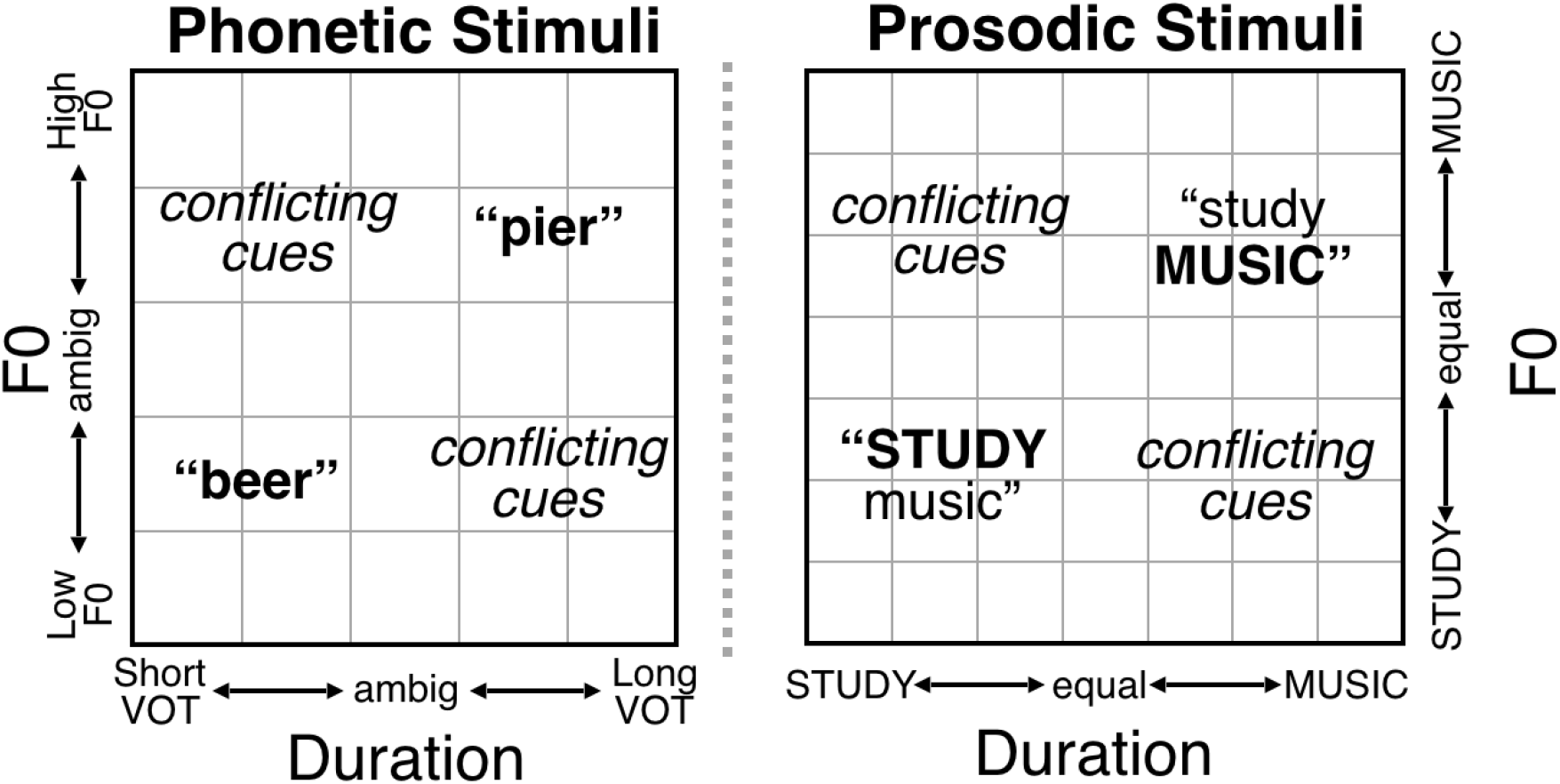
Schematic depiction of Phonetic and Prosodic stimulus spaces. In each stimulus set, a different linguistic interpretation was cued, to varying degrees, by the F0 contour and the duration of elements. Sometimes F0 and duration were indicative of the same interpretation, (upper right and bottom left corners) and other times the cues conflicted (upper left and bottom right). For the phonetic stimuli, 5 levels of an initial F0 excursion of the vowel (a pitch-related cue) and 5 levels of voice onset time (a duration cue) were crossed to create a phonetic stimulus space (Idemaru & Holt, 2011). For prosodic stimuli, 7 morphing rates of F0 and 7 of duration between “STUDY music” and “study MUSIC” were varied independently to create the stimulus space.

To ensure that the intervals along the F0 dimension were large enough that amusics would be capable of detecting them, we measured the median fundamental frequency (F0) of the words “study” and “music” in Praat (Boersma, 2002) (averaged across the entire word) for each of the seven levels of F0 morphing, then calculated the difference in F0 in semitones between the two words for each level. The average inter-level F0 differences were −8.5, −5.0, −2.1 0.6, 3.4 5.7, and 8.1 semitones. The difference in F0 between adjacent levels was always greater than 2 semitones. Since amusics are capable of detecting changes of 2 semitones and greater just as well as control participants (Hyde & Peretz, 2004; Mignault Goulet, Moreau, Robitaille, & Peretz, 2012; Moreau, Jolicœur, & Peretz, 2013; Peretz, Brattico, & Tervaniemi, 2005), and given that all of our participants had pitch discrimination thresholds of less than 1.5 semitones (see Experiment 2), we conclude that the amusic participants in this study were capable of detecting the difference between stimuli varying along the F0 dimension. As a result, this paradigm is appropriate for investigating whether amusics perceptually weight potentially informative F0 information in phonetic and prosodic categorization.

### Procedure

#### Pitch and Duration Thresholds

For all participants (Experiments 1 and 2, see below), pitch difference and duration difference thresholds were obtained in the laboratory with MLP (Grassi & Soranzo, 2009), an adaptive thresholding procedure based on the maximum likelihood method. Participants completed 3 blocks of 30 trials for both the ‘pitch’ and ‘duration’ threshold tests. On each trial participants heard 3 complex tones in a 3 alternative forced choice (3AFC) design. All complex tones comprised a fundamental frequency plus 3 harmonics, with 10ms onset and offset cosine gates. For the pitch test, participants heard 3 complex tones that were 270 ms in duration. Two of these had an F0 of 330 Hz and one had a slightly higher F0 (with F0-difference adaptively determined via the MLP procedure). The order of presentation of the higher-F0 tone was randomized (first, second or third position). Participants indicated its position by pressing keys 1, 2, or 3 on a keyboard.

For the duration test, participants heard 3 complex tones (F0 = 330 Hz). Two of these were 270 ms in duration and one was slightly longer, with the longer tone appearing randomly in any position and participants indicating its position pressing, 1, 2 or 3 on the keyboard. The threshold was calculated as the pitch or duration value that led to correct responses on 80.9% of trials on a block-by-block basis. For both the duration and pitch threshold tests, the median threshold value across the 3 blocks was extracted for statistical analysis.

#### Speech-in-noise threshold

Participants’ ability to perceive speech in the presence of background noise was also assessed; this skill is known to be impaired in tone-language speaking amusics (F. Liu, Jiang, Wang, Xu, & Patel, 2015) but has not been examined in speakers of non-tonal languages, who may rely more on duration cues when perceiving speech-in-noise (Lu & Cooke, 2009). Participants completed a speech-in-noise test, adapted from Boebinger (2015). On each trial, a participant was presented with a short sentence from the Bamford-Kowal-Bench (BKB) corpus (Bench, Kowal, & Bamford, 1979) spoken by a female talker in the presence of competing background voices (multi-talker babble). Maskers were presented at a constant level across trials while the loudness of the target voice was varied. Participants verbally reported as much of the speech as they comprehended to the experimenter, who marked how many key words were reported correctly (max of 3). The test adapted the signal-to-noise level using a one up, one down staircase procedure. The target level was varied first with an 8-dB step. The step reduced to 6-dB after the first reversal, 4-dB after the second reversal, and 2-dB after the third reversal. The procedure ended after 6 reversals at the smallest step size and the outcome measure was reported as the mean SNR level visited at the smallest step size.

#### Cue-weighting

The experiment began with instructions to listen to each item and classify whether the initial consonant was “B” or “P” (for the phonetic task) or whether the emphasis resembled “STUDY music” or “study MUSIC” (prosodic task). Responses were made by clicking a mouse to indicate one of two buttons positioned near the center of the screen (“B”/”STUDY music” on left; “P”/”study MUSIC” on right). Each phonetic block contained 50 trials (2 measurements at each of the 25 combinations of F0 and duration). Each focus block contained 49 trials (1 measurement at each of the 49 F0 and duration combinations). Each participant completed 20 blocks in total: 10 phonetic blocks (500 trials total, 20 repetitions of each stimulus in the phonetic acoustic space) and 10 prosodic blocks (490 trials total, 10 repetitions of each stimulus in the prosodic acoustic space). Blocks alternated between phonetic and prosodic tasks, with short breaks interspersed. The entire experiment lasted approximately 60 minutes. Participants completed the experiment online, at home in a quiet room alone, using either headphones or external speakers (which was confirmed by email after testing). Two subjects did not have computers with sound capability and were therefore tested in a laboratory at Birkbeck College. For all participants, testing was conducted via the Gorilla web experiment platform (www.gorilla.sc/about) (Anwyl-Irvine, Massonnié, Flitton, Kirkham, & Evershed, 2018).

#### Statistical analysis

Data were analyzed with R (R Core Team 2018). Cue weights were calculated by constructing a multiple logistic regression for each participant (separately for phonetic and prosodic components) with F0 and Duration as factors (on integer scales from 1-5 for the phonetic task and 1-7 for the prosodic task, according to the number of stimulus increments). The coefficients estimated from these models were then normalized such that the F0 and Duration weights summed to one to indicate the relative perceptual weight of each acoustic dimension in categorization responses (Holt & Lotto, 2006; Idemaru et al., 2012). A large coefficient for F0 relative to Duration indicated that the F0 factor explained more variance in participants’ categorization judgments than the Duration factor. To test for group effects, the Cue Weights for F0 and Duration were extracted for each participant and subjected to an independent-samples t-test, separately for phonetic and prosodic tasks. Because the distributions of pitch thresholds were non-normal, the relationships between pitch and duration thresholds and cue weights were tested with Kendall’s Tau-b.

### Results

#### Pitch, Duration and Speech-In-Noise Thresholds

Among the subset of participants who took part in Experiment 1, amusic participants, compared to controls, had significantly higher pitch thresholds (Wilcoxon Rank Sum W = 107, p=.002), but similar duration (W=50, p=.51) and speech-in-noise thresholds (W = 78.5, p=.25). The pattern held across all participants: amusics as a group had higher pitch thresholds than controls (W = 29, p < 0.001), but did not differ from controls in tone duration discrimination (W = 129, p = 0.74) or speech-in-noise threshold (W = 155.5, p = 0.17; Fig. 2). For the amusic group, all pitch discrimination thresholds were below 1.5 semitones, assuring that our manipulation of the F0 dimension in 2-semitone steps in the cue weighting paradigms would be perceptible even to amusic participants.

**Figure 2:**
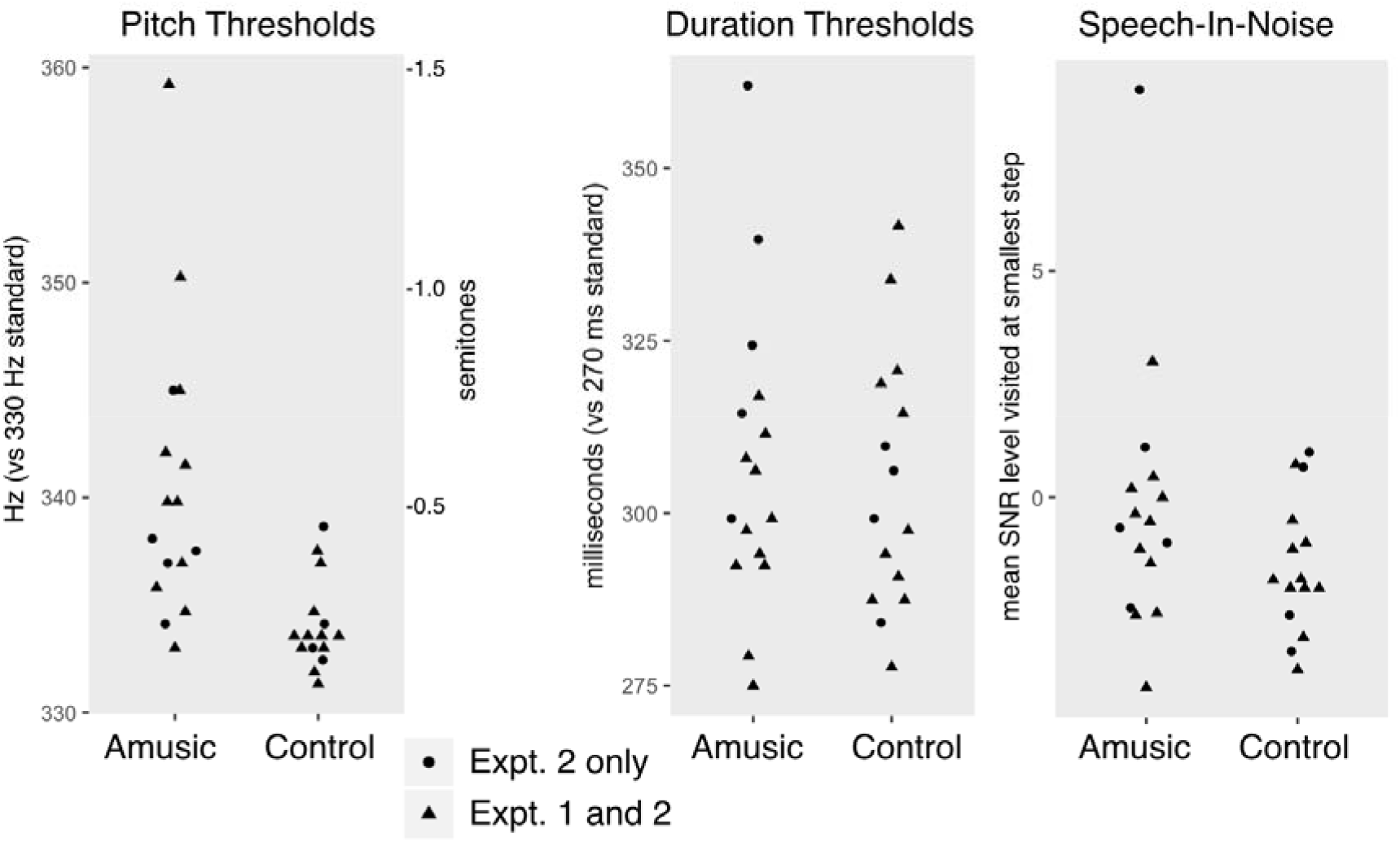
Thresholds for detecting pitch changes, duration changes, and for speech-in-noise.

#### Phonetic Cue Weighting

The normalized cue weights for the amusic and control groups were compared with t-tests because normalizing the F0 and duration cue weights relative to each other (so they sum to 1) causes them to be non-independent, and therefore a Group X CueType interaction test was inappropriate. Recall that duration (VOT) was expected to be the primary cue for the phonetic task. Accordingly, the magnitude of the pitch and duration weights did not differ between groups (Fig. 3A; group effects T_(20)_= 1.58, *p* = 0.13). Although the mean responses between amusics and controls (Fig. 4A) did not differ, the results of the independent-samples t-test of perceptual cue weights across groups across each of the 25 stimuli are presented in Fig. 4B for illustrative comparison. Here, again, categorization performance across groups was similar for each stimulus in the acoustic space. No group differences were detected even at a very lenient (uncorrected for multiple comparisons) threshold of *p* < 0.05.

**Figure 3:**
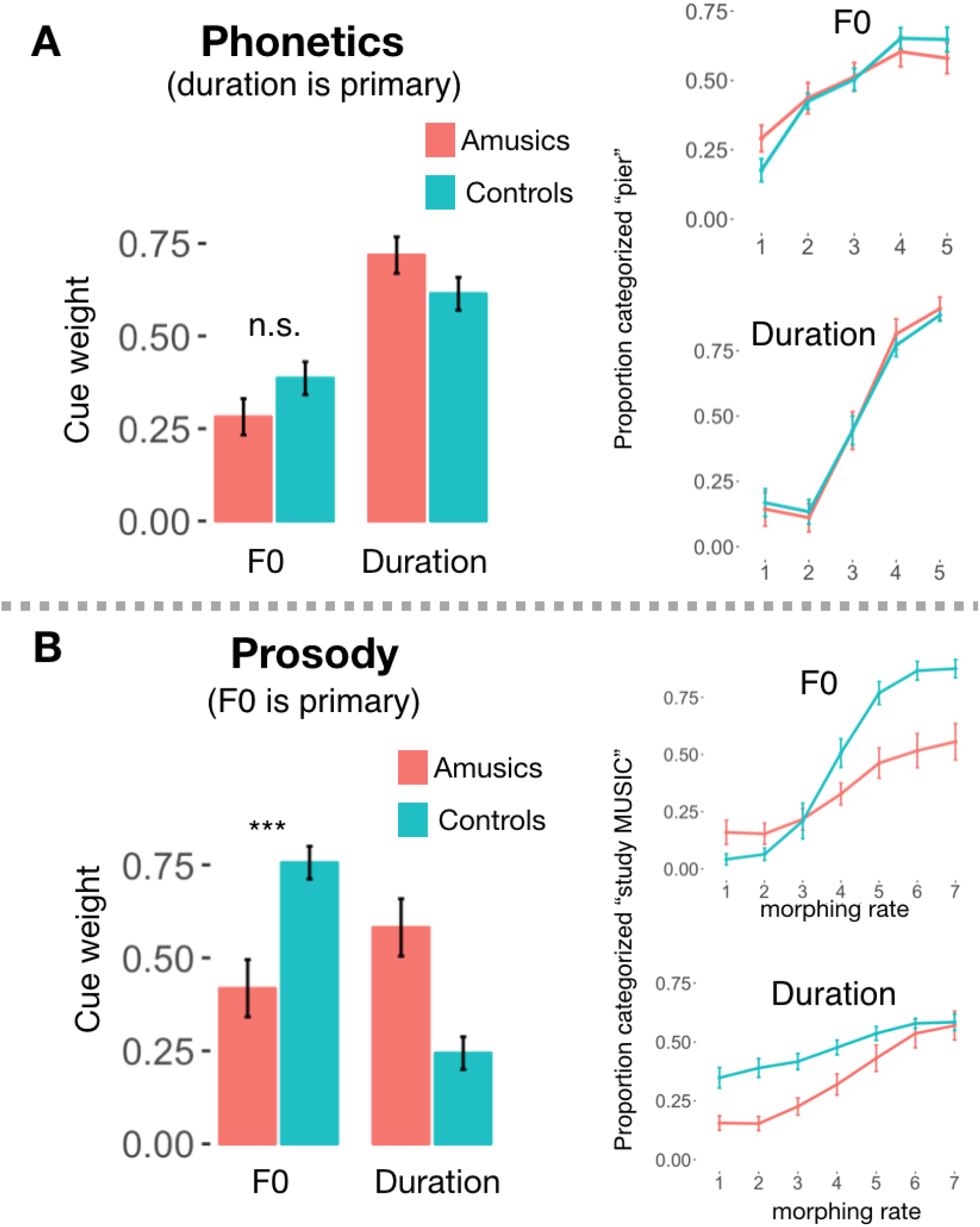
Comparison of F0 and duration cue weights for phonetic and prosodic perception. A) Phonetic Cue Weighting. Mean perceptual weights plotted by group and condition (left). Mean categorization response plotted at each level of F0, collapsed over duration; and each level of duration collapsed over F0. **B) Prosodic Cue Weighting.** Analogous plots for the prosodic categorization. Bars indicate standard error of the mean.

**Figure 4:**
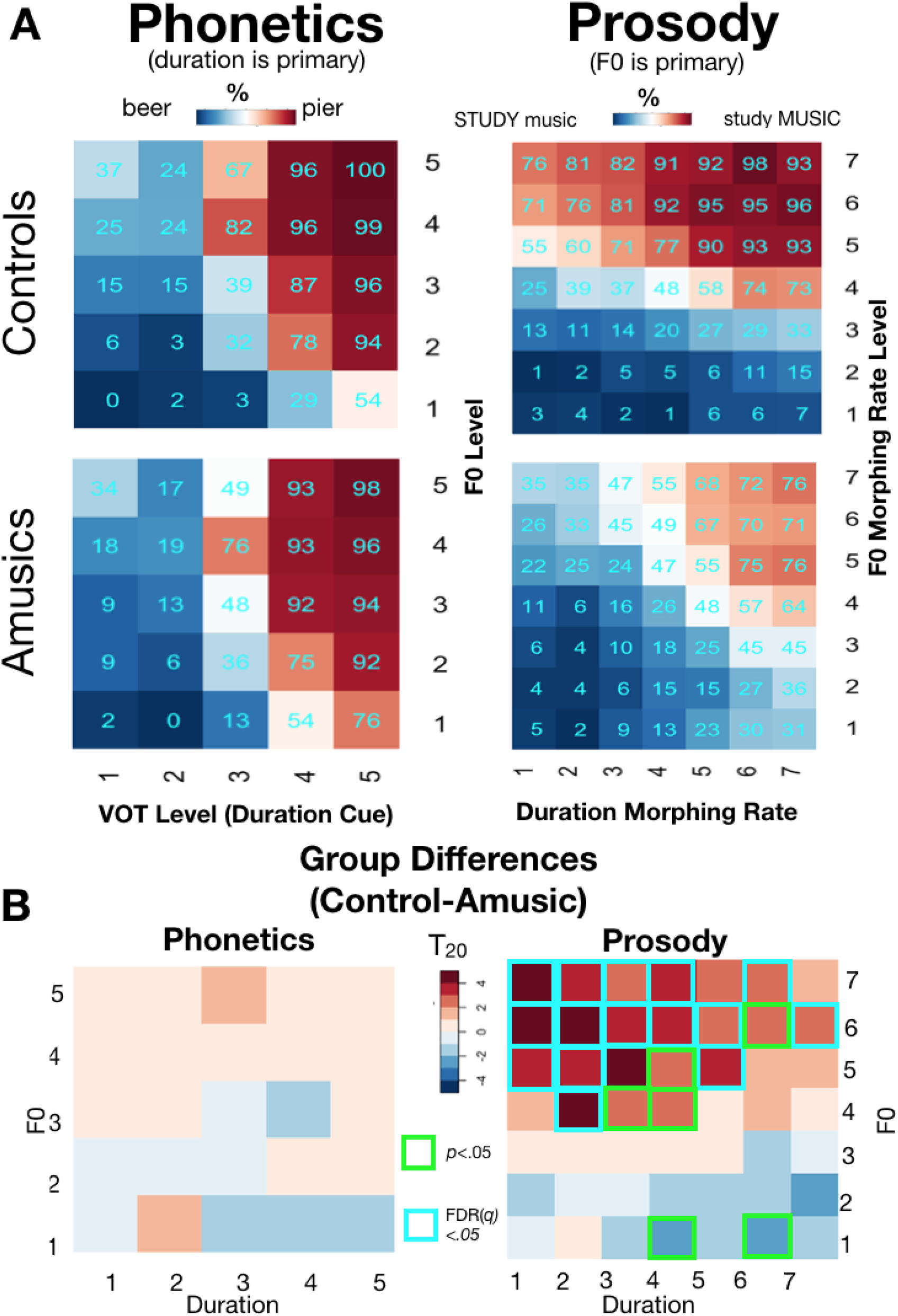
When F0 and duration cues conflict, amusics rely on duration when categorizing prosody. **A)** Heatmaps indicate proportion of trials categorized as “study MUSIC” (for the prosody portion, panel A) or “pier” (phonetics portion, panel B), for the Control and Amusic groups. **B)** Group difference (Control – Amusic) heatmaps displaying T-statistics. When duration and F0 conflicted in the prosody task (duration indicated emphasis on STUDY, but F0 indicated emphasis on MUSIC; upper-left quadrant of stimulus space), amusic participants chose the duration-based response more often than controls. Teal outlines indicate significant group differences (corrected for multiple comparisons). Uncorrected results (*p*<.05) are indicated with green outlines.

Finally, we calculated a standard score reflecting participants’ relative ability to discriminate pitch and duration in complex tones based on their perceptual thresholds, and tested whether this metric was related to subjects’ relative perceptual weightings in the phonetics task. To obtain a standard score each subject’s pitch and duration threshold were subtracted from the standard F0 (for pitch, 330 Hz) or standard duration (270 ms) used in the psychophysics test, then divided by the standard deviations of these distributions across subjects. The standard scores for pitch and duration were then combined with an asymmetry ratio [(Duration – Pitch) / (Duration + Pitch)] such that higher values indicated finer pitch than duration thresholds and lower values indicated the reverse. The relative ability to discriminate tone pitch versus duration did not correlate with perceptual weight of F0 in phonetic categorization (Kendall Tau-b *r*=-0.12, *p*=.45; Fig. 5A). Full results for each individual subject, for both the phonetics and prosody tasks are plotted in Figure 6.

**Figure 5:**
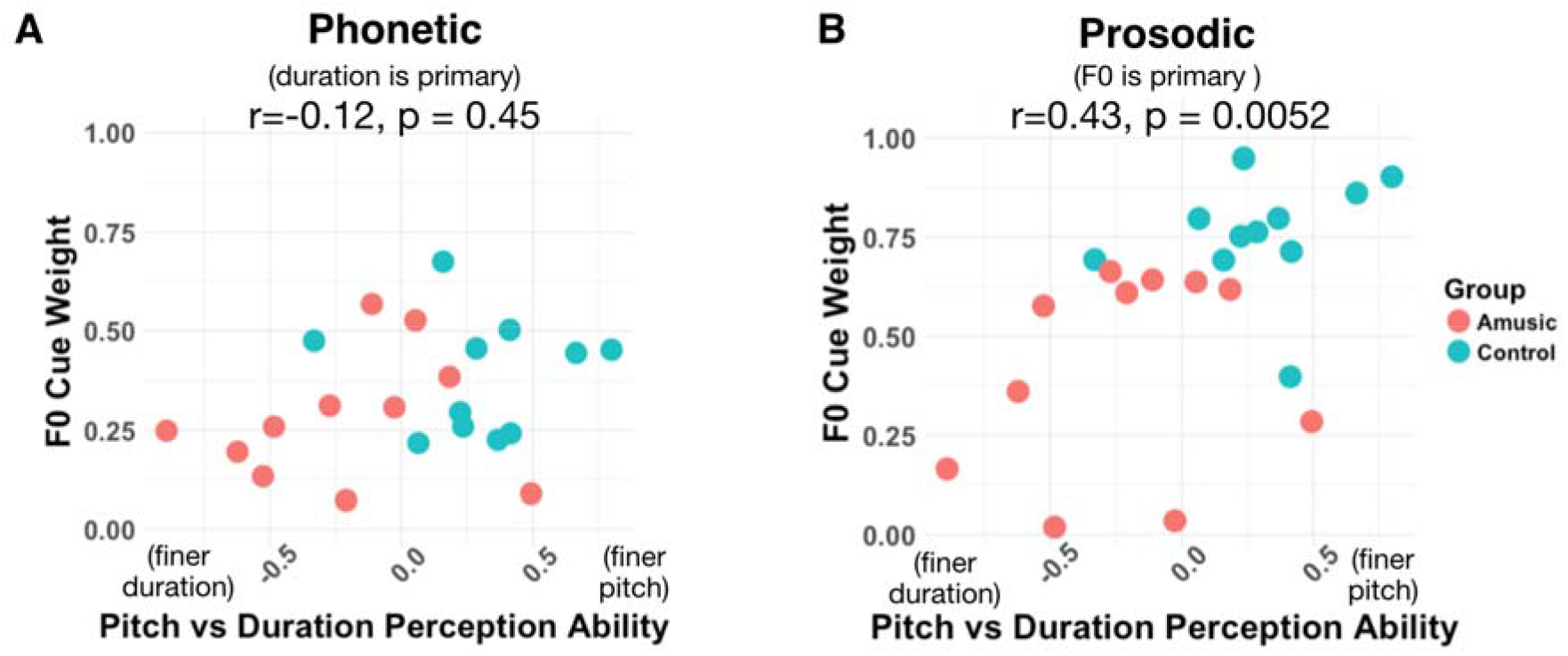
Correlation between perceptual discrimination and perceptual cue weight. The figures plot relative pitch versus duration thresholds as a function of the relative perceptual weight of F0 in phonetic (A) and prosodic (B) judgments. The correlation shown is Kendall’s Tau-b. **A.** In the phonetic task, for which F0 is a secondary perceptual dimension, there was no significant correlation between perceptual thresholds and normalized perceptual weight of F0. **B.** In the prosodic task, for which F0 is the primary perceptual dimension, individuals with finer pitch discrimination thresholds tended to rely more on F0 in prosodic focus judgements, even though the F0 differences in the prosodic task were large enough to be perceptible by all participants.

**Figure 6:**
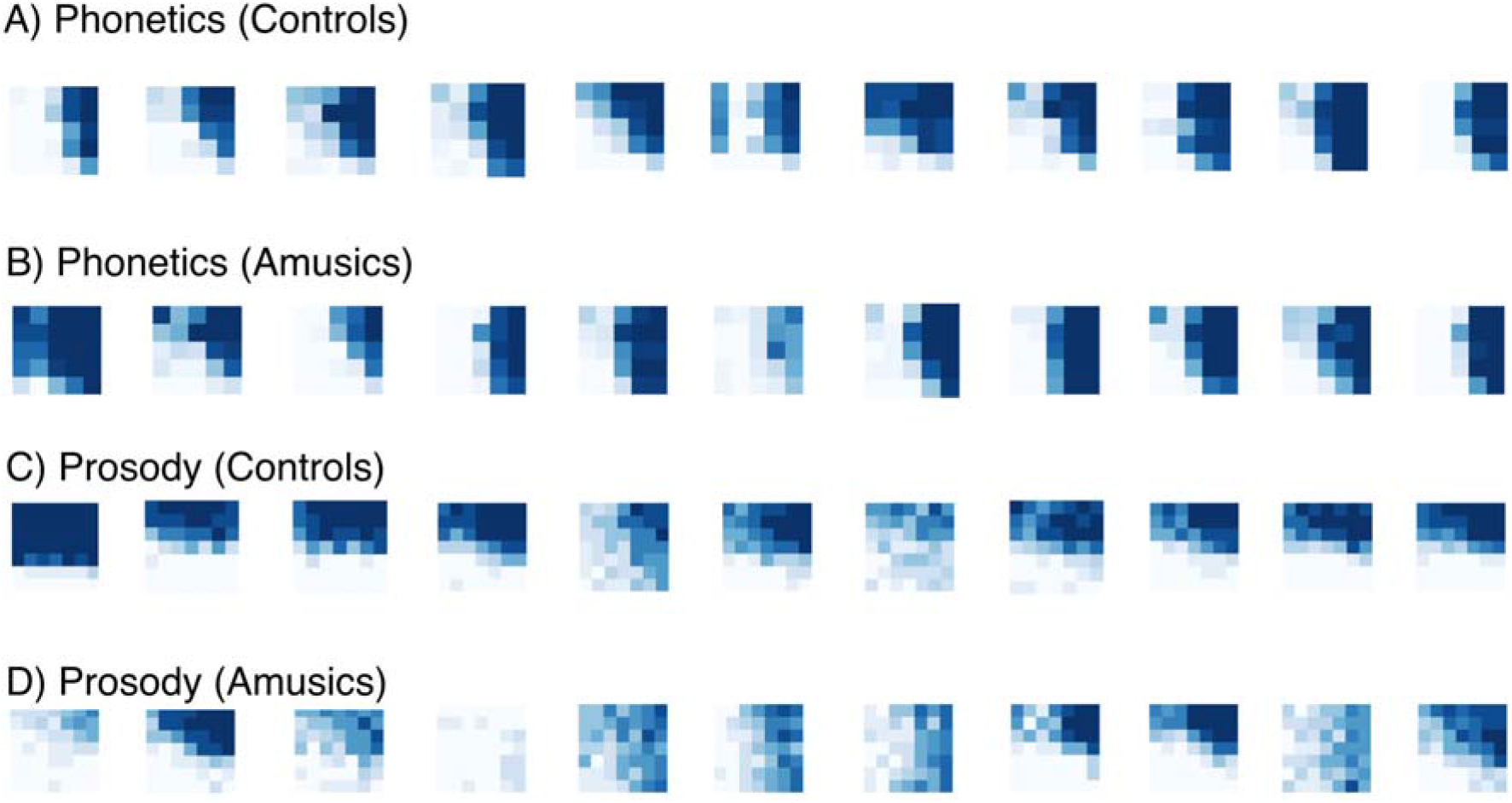
Individual heatmaps indicating categorization responses. Categorization responses for Experiment 1 subjects for the prosody blocks (A-B) and phonetic blocks (C-D). Horizontal axis indicates duration and vertical axis indicates F0. Plots in A,C and in B,D present the participants in the same order (e.g. data from leftmost plot in A is from the same participant as leftmost plot in C).

#### Prosodic Cue Weighting

In the prosodic cue weighting task, for which F0 was the primary cue, the results were strikingly different. The relative F0 perceptual weights were higher in controls than amusics, whereas duration weights were higher for amusics than controls (both group comparisons T_(20)_ = 3.81, *p*=.001, Fig. 3B). This suggests that amusics, who do not process pitch reliably in general, indeed perceptually weighted duration over F0. As mentioned above, an interaction test on the normalized (relative) cue weights is inappropriate because they sum to 1 (e.g. if a participant’s pitch weight were 0, his or her duration weight would necessarily be 1); however, we note that an ANOVA on the raw cue weights (before normalization) further confirmed this pattern (interaction of CueType [pitch vs duration] X Group [Amusic vs Control], F_(1,39)_ = 10.3, *p* = 0.002).

To provide a finer-grained understanding of each group’s weighting, Figure 4A plots mean responses for each group across each of the 49 stimuli depicted schematically in Figure 1. As in the phonetic cue weighting task, we compared these matrices cell-by-cell (two-sample T-tests Controls>Amusic, FDR-corrected, Fig. 4B). Unlike in the phonetic task for which Control and Amusic participants performed similarly, many group differences emerged in the prosodic categorization task, where controls relied relatively more on pitch versus duration compared to amusics. All (multiple-comparisons-corrected) significant differences were in the top half of the matrix, where F0 cued an emphasis on “MUSIC”. Most (12 out of 16) of these significant group differences occurred in the 16 stimuli of the upper-left quadrant, where emphasis was placed on “STUDY” by duration cues, and “MUSIC” by F0 cues. In this ambiguous quadrant of the stimulus space, where F0 and duration signaled differing interpretations, amusics relied on the duration cues to make their response more often than did controls.

Across both the amusic and control groups, participants with finer pitch than duration discrimination thresholds tended to have higher F0 (than duration) cue weights (Kendall Tau-b *r*=0.43, *p*=.005; Fig. 5B).

Finally, we confirmed statistically the greater group differences in perceptual weightings in the Prosody task compared to the Phonetic task (interaction of Group X Experiment predicting normalized F0 cue weights, F_(1,40)_ = 4.44, *p*=.04).

## Discussion

In two categorization tasks, participants with amusia and control participants categorized stimuli varying across F0 and duration dimensions according to a phonetic distinction and a prosodic distinction. The phonetic and prosodic categorization tasks varied in the extent to which category membership was cued by information from the F0 dimension versus the duration dimension. For the phonetic distinction, duration had been established by prior literature to be the primary dimension while for the prosodic distinction, F0 was known to be primary (Haggard et al., 1970; Massaro & Cohen, 1976; 1977). We found that when duration was the primary dimension (in the phonetics task), the amusic and control groups categorized similarly, each relying predominantly on duration (VOT) to signal category membership. However, when F0 was primary (in the prosody task), classification behavior differed between the groups: controls classified ambiguous stimuli (for the most part) according to the primary dimension (F0), as predicted by prior research (Breen et al., 2010) whereas amusics tended to classify the very same stimuli instead according to duration. Crucially, this difference could not be explained by the amusic participants being simply unable to hear the F0 cues, because (1) sensitivity to the F0 dimension was observed in both the phonetic and prosodic task and (2) the differences in F0 across stimulus levels in each task were large enough to be above-threshold and clearly perceptible to the amusics. Indeed, they were well above amusics’ independently-measured pitch thresholds.

These results suggest that amusic individuals have developed perceptual strategies, focusing on the dimensions which they are best able to process and to the exclusion of the dimensions which they have historically found less useful. This could help explain why amusics do not generally report difficulty with speech in everyday life (F. Liu et al., 2010), despite displaying deficits when tested on speech perception tasks requiring the use of F0 information. Indeed, we further found that psychophysical thresholds across all participants, controls and amusics, correlated with the relative weighting of F0 versus durational information in the prosodic task, confirming a link between the precision of perception along a given primary dimension and the extent to which listeners weight that dimension in perceptual categorization. This relationship between perceptual acuity and categorization strategies could extend to other perceptual domains as well, such as identification of characteristics of environmental sounds (Lutfi & Liu, 2007).

The results of Experiment 1 revealed that amusics weight F0 cues relatively less than duration cues, compared to controls. This was true for the prosodic task where F0 was typically the primary dimension, but not true for the phonetic task, where duration is typically primary. This suggests that listeners will only change cue weighting in response to perceptual difficulties if the dimension they have difficulty processing is the primary cue for a given linguistic feature. An alternative explanation, however, is that these findings may reflect a specific amusic deficit for integration of pitch information across longer time frames, rather than a simple encoding deficit. Indeed, in the prosody paradigm, F0 information unfolded over time in the order of seconds, whereas in the phonetic paradigm the F0 excursion was over milliseconds. Future work could distinguish between these two explanations (primary vs secondary cue; long vs. short time scale) by examining cue weighting in amusics and controls for perception of a prosodic feature for which duration is a dominant cue, such as lexical stress (Mattys, 2000).Overall, these results indicate that the manner in which auditory dimensions map to linguistically-relevant categories differs across listeners, and that these individual differences in perceptual weighting reflect, at least in part, variability in how well listeners can perceive auditory dimensions. A particular dimension, for example, may be primary for one listener but secondary for another, leading to differing categorization strategies for the same stimuli across different listeners.

## Experiment 2

In Experiment 2 we examined whether cue redundancy in more naturalistic speech and music stimuli leads to robustness in the face of perceptual deficits by examining music and speech perception in amusics and control participants. In other words, we tested whether the ‘prosodic strategy’ revealed by the results of Experiment 1—where amusics rely less on pitch information when it is a primary cue, focusing instead on other sources of information about speech—can be successful enough to lead to perceptual categorization performance equivalent to that of control participants.

First, although amusia is canonically a problem with using pitch information to perceive music, there are temporal cues to certain musical structures (such as the beginnings and ends of phrases) that could theoretically enable amusics to compensate for poor pitch perception and successfully track musical features. To examine this possibility, in a music perception test, participants categorized phrases as complete or incomplete when they could rely on pitch alone, duration alone, or both dimensions simultaneously. Second, we investigated whether temporal dimensions in speech provide sufficient information for amusics to compensate for poor pitch perception, enabling categorization of prosodic information at a level similar to that of controls. In two linguistic prosody tests we measured the extent to which participants could use F0 and duration dimensions to make categorical decisions about whether words had linguistic focus (or ‘emphasis’; e.g. ‘Mary likes to READ books, but she doesn’t like to WRITE them.’) or whether phrase boundaries were present or absent (’After John runs [phrase boundary], the race is over’). To do this, we manipulated stimuli such that participants needed to rely on the F0 dimension alone, the duration dimension alone, or could use both. Across both speech and music perception, if amusics rely more on durational dimensions (for which they are putatively unimpaired), performance should be more accurate when they can rely on redundant information (F0 and duration), compared to when they must rely solely on a dimension that is less reliably perceived (F0). Indeed, we predicted that in the presence of durational information amusics would be unimpaired relative to control participants, even for the detection of musical features; that is, duration cues may ‘bootstrap’ perception that would ordinarily be driven by F0 and pitch contours.

## Methods

### Participants

For description of participants see Experiment 1.

### Musical Phrase Perception Test

#### Stimuli

The stimuli consisted of 100 musical phrases taken from a corpus of folk songs (Schaffrath & Park, 1995). Musical phrases were synthesized as a sequence of six-harmonic complex tones with 10ms cosine onset and offset ramps. Stimuli were created in three conditions that manipulated the acoustic dimensions available to listeners. In the Combined condition, the musical phrase contained both pitch and duration information (as typical naturalistic melodies do). Fifty stimuli were selected from the database for this condition only, and fifty additional stimuli were selected for each of two additional conditions. In the Pitch condition, the pitch of the melody was preserved (as in the original version) but the durations were set to be isochronous and equal to the mean duration of the notes in the original melody. In the Duration condition, the original note durations were preserved but the pitch of the notes was made to be monotone at a pitch of 220 Hz. In an additional manipulation, half of the stimuli presented in each condition formed a complete musical phrase with the notes in an unmodified sequential order - these could be perceived as a *Complete* musical phrase. The other half were made to sound *Incomplete* by presenting a concatenation of the second half of the musical phase and the first half of the next musical phrase in the song. The order of the notes within the two halves was preserved. Thus, the resulting *Incomplete* stimuli contained a musical phrase boundary that occurred in the middle of the sequence, rather than at the end.

#### Procedure

On each trial, a musical note sequence was presented to the participant through headphones. After the sound finished playing, a response bar appeared on the screen which was approximately 10 cm in width. Subjects were tasked with deciding how complete each musical phrase sounded by clicking with their mouse on the response bar. (A continuous measure of completeness was used to avoid potential ceiling/floor effects due to bias on the part of individual listeners to hear phrases as complete or incomplete.) The word “Incomplete” was shown on the left side of the response bar, and the word “Complete” was shown on the right. Participants could click anywhere within the bar to indicate how complete they thought the phrase had sounded (Fig. 7). The next stimulus was played immediately after a response. Participants judged 3 blocks of 50 trials each, with a short break in between. As the study was aimed at understanding individual differences, the block order was always the same, with all the trials in a condition presented in a single block (Combined Cues, then Duration Only, then Pitch Only). Within a block, half of the trials were Complete, and half Incomplete. The main outcome measure was the raw rating difference between Complete and Incomplete trials for each condition.

**Figure 7.**
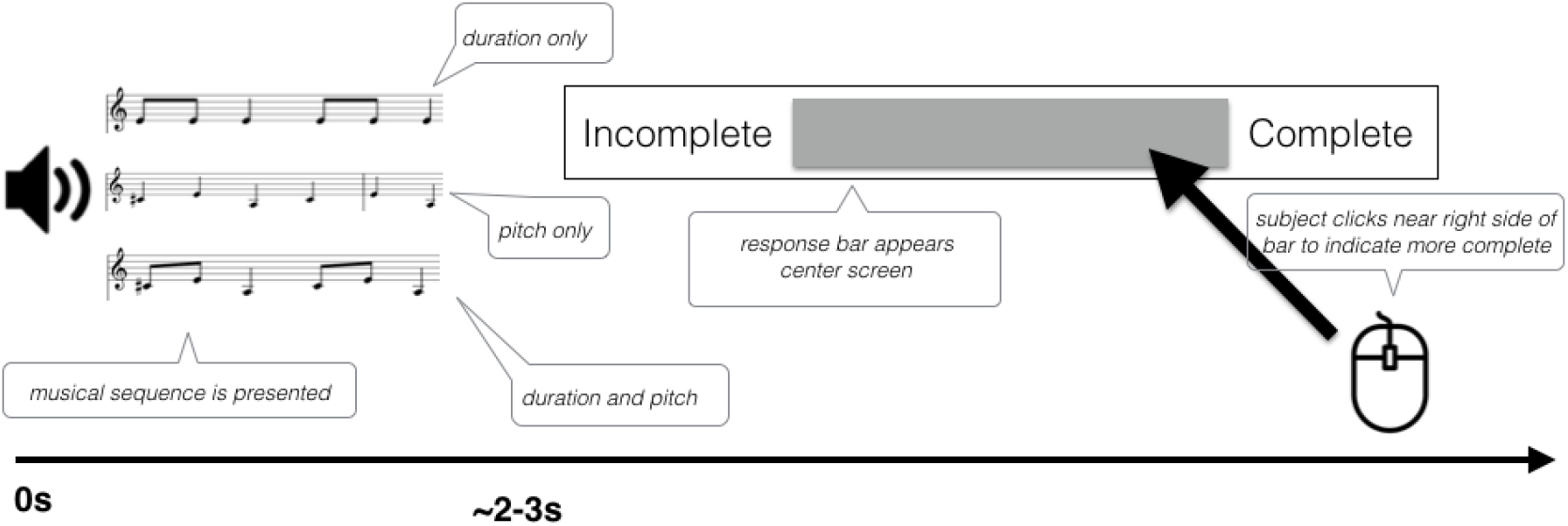
Schematic of trial structure for the Musical Phrase Test. Participants heard either a complete musical phrase or a musical sequence that straddled the boundary in between two musical phrases. They then indicated how complete they thought the phrase sounded by clicking with a mouse at a point along a response bar.

### Linguistic focus perception task

#### Stimuli

The stimuli consisted of 47 compound sentences with an intervening conjunction created specifically for this study. Each of the sentences was read with emphasis to create had two versions: “early focus”, where emphasis occurred early in the sentence and served to contrast with a similar word later in the sentence (e.g., “<u>Mary likes to READ books</u>, but she doesn’t like to WRITE them,” focus indicated by upper-case letters), and “late focus”, where the focus occurred slightly later in the sentence (e.g., “<u>Mary likes to read BOOKS</u>, but she doesn’t like to read MAGAZINES,” focus indicated by upper-case letters). (See Fig 8A,B; Supplemental Appendix 1).

**Figure 8.**
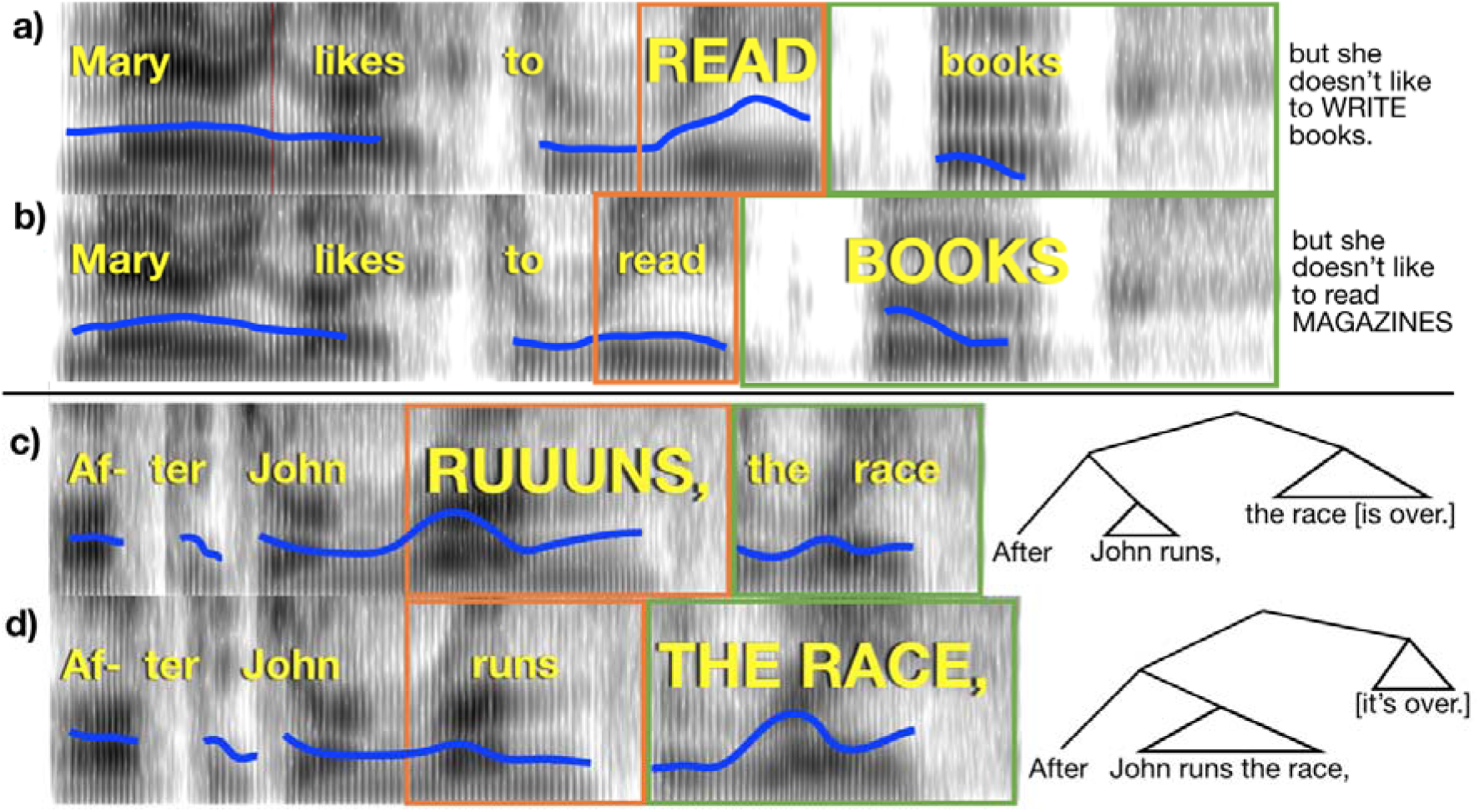
Pitch and duration correlates of emphatic accents and phrase boundaries. Spectrograms of stimuli used in the experiment (time on horizontal axis, frequency on vertical axis, and amplitude in grayscale), with linguistic features cued simultaneously by F0 and duration (the “Combined” condition). Blue line indicates F0 contour. Width of orange and green boxes indicate duration of the words within the box. A) Emphatic accent places focus on “read”. Completion of the sentence appears to the right. B) Emphatic accent places focus on “books”; sentence completion is at right. C) A phrase boundary occurs after “runs”. D) A phrase boundary occurs after “race”. Syntactic trees are indicated at right to illustrate the structure conveyed by acoustics of the stimuli.

We recorded these sentences (44.1 kHz, 32-bit) using a Rode NT1-A condenser microphone as they were spoken by an actor who placed contrastive accents to emphasize the capitalized words. Recordings of both versions of the sentence were obtained and cropped to the identical portions (underlined above). Using STRAIGHT software (Kawahara & Irino, 2005), the two versions were manually time aligned by marking corresponding anchor points in both recordings. We then produced 6 different kinds of morphs by varying the amount of pitch-related (F0) and duration information either independently or simultaneously. For *F0 only* stimuli pairs, the late and early focus sentences differed only in F0 excursions. The temporal morphing proportion between the two versions was held at 50% while the F0 was set to include 75% of the early focus version or 75% of the late focus version recording. This resulted in two new ‘recordings’ that differed in F0 excursions, but were otherwise identical in terms of duration, amplitude and spectral quality. For *duration only* stimuli, we created two more morphs that held the F0 morphing proportion at 50% while the duration proportion was set to either 75% early focus or 75% late focus. The output differed only in duration, and were identical in terms of F0, amplitude and spectral quality. Finally, we made *“naturalistic”* stimuli where both F0 and duration information contained 75% of one morph or the other, and thus F0 and duration simultaneously cued either an early or late focus reading. All files were saved and subsequently presented at a sampling rate 44.1 kHz with 16-bit quantization.

#### Procedure

Stimuli were presented with Psychtoolbox in Matlab. Participants saw sentences presented visually on the screen one at a time, which were either early or late focus (see paradigm schematic in Fig 8A,B and Fig 9A). The emphasized words appeared in all upper-case letters, as in the examples above. Subjects had 4 seconds to read the sentence to themselves silently and imagine how it should sound if someone spoke it aloud. Following this, subjects heard the first part of the sentence spoken aloud in two different ways, one that cued an early focus reading and another that cued late focus. Participants were instructed to listen and decide which of the two readings contained emphasis placed on the same word as in the text sentence. After the recordings finished, subjects responded by pressing “1” or “2” on the keyboard to indicate if they thought the first version or second version was spoken in a way that better matched the on-screen version of the sentence. The correct choice was cued either by F0 or duration exclusively, or both together. The serial order of the sound file presentation was randomized. The stimuli were divided into 3 lists (47 trials each) and counterbalanced such that each stimulus appeared once in each condition (F0, duration, and both). For half (23) of the stimuli, two of the presentations were early focus, and one was late focus; for the remaining stimuli, two presentations were late focus and one was early. The entire task lasted approximately 30 minutes.

**Figure 9:**
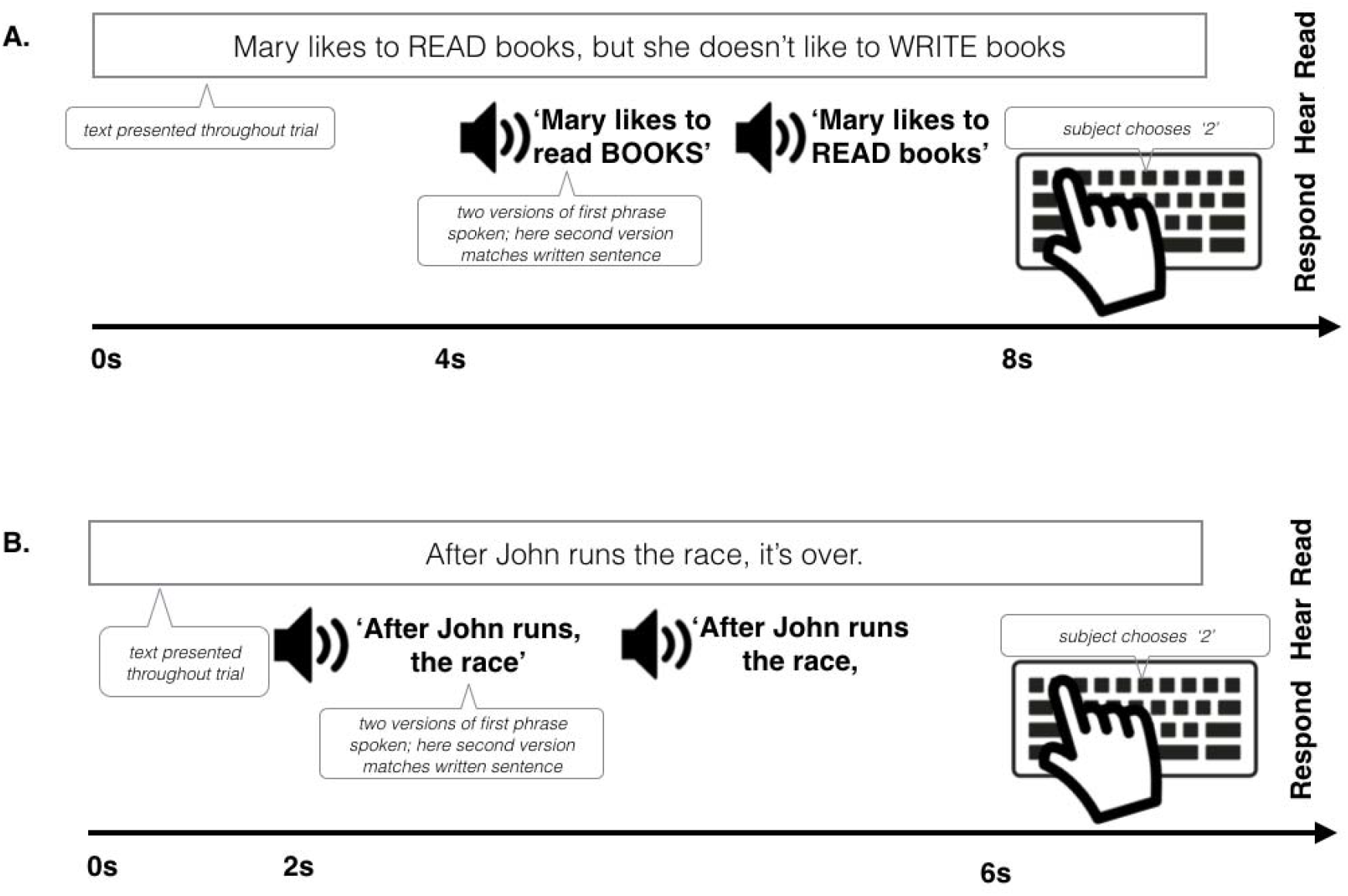
Example trial structure for the linguistic focus test (A) and the linguistic phrase test (B). First, a single sentence was presented visually, and the participants read it to themselves. Next, two auditory versions of the first part of the sentence were played sequentially, only one of which matched the focus pattern of the visually presented sentence. Participants then indicated which auditory version matched the onscreen version with a button press.

### Linguistic phrase perception test

#### Stimuli

The stimuli consisted of 42 short sentences with a subordinate clause appearing before a main clause (see Figure 8C, D). About half of these came from a published study (Kjelgaard & Speer, 1999) and the rest were created for this test (see Supplemental Appendix 2). The sentences appeared in two conditions: an “early closure” condition, for which the subordinate clause’s verb was used intransitively, and the following noun was the subject of a new clause (“After John runs, the race is over”); and “late closure”, where the verb was transitive and took the following noun as its object, causing the phrase boundary to occur slightly later in the sentence (“After John runs the race, it’s over”). Both versions of the sentence were lexically identical from the start of the sentence until the end of the second noun.

A native Standard Southern British English-speaking male (trained as an actor; the same voice model as for the focus task) recorded early and late closure versions of the sentences in his own standard Southern English dialect. The recordings were cropped such that only the lexically identical portions of the two versions remained, and silent pauses after phrase breaks were excised. The same morphing proportions were used as in the focus task – with early or late closure cued by 75% morphs biased with pitch, duration or both combined. As was done with the linguistic focus task, the stimuli were crossed with condition (F0, duration, and both) and early vs. late closure and divided into three counterbalanced lists.

#### Procedure

The procedure for the Linguistic Phrase test was similar to that of the Linguistic Focus Test. Participants saw sentences presented visually on the screen one at a time, which were either early or late closure, as indicated by the grammar of the sentence and a comma placed after the first clause (Figure 9B). They then had two seconds to read the sentence to themselves silently and imagine how it should sound if someone spoke it aloud. Following this period, subjects heard the first part of the sentence (which was identical in the early and late closure versions) spoken aloud, in two different ways, one that cued an early closure reading and another that cued late closure. The grammatical difference between the two spoken utterances on each trial was cued by either F0 differences, duration differences, or both F0 and duration differences. Subjects completed three blocks of 42 trials. Stimuli were counterbalanced such that each stimulus appeared once in each condition, and half the of presentations were early close and half were late close. The task was performed in a lab at Birkbeck and lasted approximately 25 minutes.

### Psychophysics

As described in Experiment 1, thresholds for pitch and duration discrimination, and ability to hear speech in noise, were measured using the MLP toolbox (Grassi & Soranzo, 2009).

### Statistical analysis

Data were analyzed with R. For the musical phrase, linguistic focus and linguistic phrase tests, linear mixed effects models were estimated using *lme4*, with Group (Amusic or Control), Condition (Pitch, Duration or Combined) and their interaction entered as fixed effects, and Item and Subject as random intercepts. P-values for these effects were calculated with likelihood ratio tests of the full model against a null model without the variable in question. Comparisons of predicted marginal means were performed with *lsmeans.*

The dependent variable for the Musical Phrase Test was calculated by identifying the raw response value between −50 and 50 (for each trial) based on the position along the response bar on which the participant clicked, with −50 corresponding to responses on the extreme end of the Incomplete side of the scale. The sign of the data point for Incomplete trials was then inverted so that more positive scores always indicated correct performance and greater scores indicated more accurate categorization of musical phrases.

The dependent variable that was entered into the model for the Focus and Linguistic Phrase tests was whether each response was CORRECT or INCORRECT. Because the dependent variable was binary, we used the generalized linear mixed models (*glmm)* function in the *lmer* package to estimate mixed effects logistic regressions, and we report odds ratios as a measure of effect size.

## Results

### Musical phrase perception

To recap, the musical phrase perception test (Fig. 7) tested participants’ ability to perceive the extent to which a series of notes resembled a complete musical phrase, with acoustic dimensions of F0, duration, or their combination available to support the judgment. The availability of acoustic dimensions affected participants’ accuracy in identifying complete versus incomplete phrases. Accuracy was highest when both cues were present, lowest when only the F0 dimension was present, and intermediate when only the duration dimension was present (main effect of Condition χ^2^(4) = 30.76, p < .001, see Table 1 for pairwise statistics). Compared to controls, amusics were overall less accurate (main effect of Group χ^2^(3) = 9.43, p = 0.02) and also differentially affected by which dimensions were present (Group x Condition interaction χ^2^(2) = 8.21, p = 0.02). FDR-corrected pairwise tests showed that when only the pitch-relevant F0 dimension was available, the average amusic’s performance was significantly lower than controls (*p*=.024; Table 1). Indeed, the confidence interval around amusics’ mean accuracy included zero (Fig. 10A; Table 1), suggesting that they were unable to perform the task using pitch cues alone. By contrast, when amusics could rely on duration alone, or both pitch and duration together (as in the Combined condition, where cues were present as in naturalistic melodies), amusics and controls did not differ significantly.

**Table 1:** Musical Phrase Test, all pairwise contrasts (p-values FDR-adjusted)

| **Condition** | **Group** | **Contrast** | **Beta** | **SE** | **df** | **T** | **p** |
| --- | --- | --- | --- | --- | --- | --- | --- |
| Combined | ~ | CONT vs AMUS | 0.7 | 1.7 | 124.78 | 0.41 | 0.683 |
| Duration | ~ | CONT vs AMUS | -1.41 | 1.7 | 124.78 | -0.83 | 0.528 |
| Pitch | ~ | CONT vs AMUS | 4.6 | 1.7 | **124.78** | **2.7** | **0.024** |
| ~ | CONT | Combined vs Duration | 3.94 | 2.82 | 142.82 | 1.4 | 0.298 |
| ~ | CONT | Combined vs Pitch | 5.42 | 2.82 | 142.82 | 1.92 | 0.128 |
| ~ | CONT | Duration vs Pitch | 1.48 | 1.53 | 4513.25 | 0.97 | 0.5 |
| ~ | AMUS | Combined vs Duration | 1.84 | 2.8 | 137.67 | 0.66 | 0.577 |
| ~ | AMUS | Combined vs Pitch | 9.32 | 2.8 | **137.67** | **3.33** | **0.005** |
| ~ | AMUS | Duration vs Pitch | 7.48 | 1.48 | **4513.25** | **5.06** | **<.001** |
| ~ | ALL | Combined vs Duration | 2.89 | 2.60 | 103.03 | 1.11 | 0.270 |
| ~ | ALL | Combined vs Pitch | 7.37 | 2.60 | **103.03** | **2.83** | **0.008** |
| ~ | ALL | Duration vs Pitch | 4.48 | 1.06 | **4513.25** | **4.21** | **<.001** |

**Figure 10.**
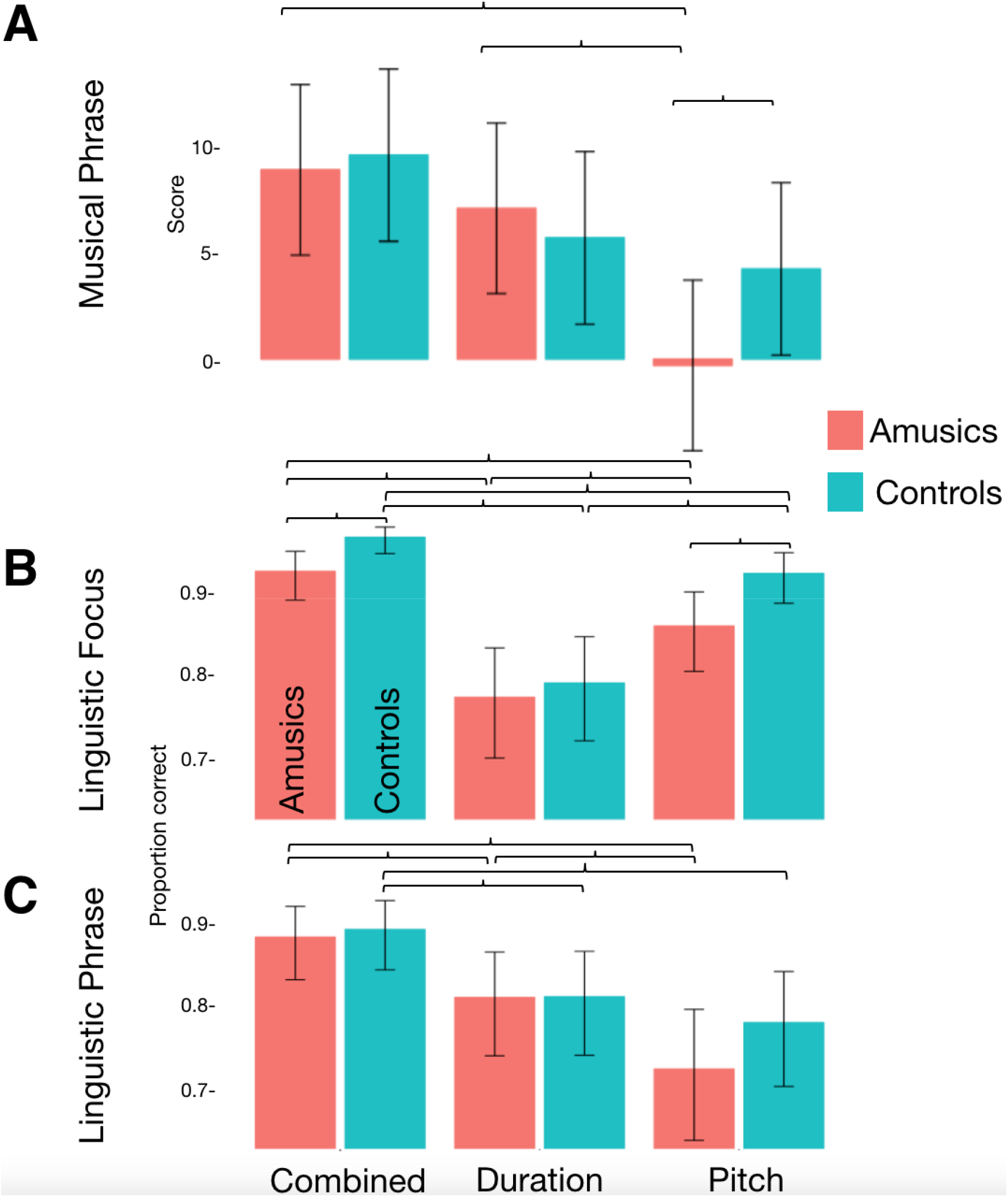
Results of the Linguistic Focus, Linguistic Phrase and Musical Phrase tests. Bars show 95% confidence intervals and brackets indicate significant pairwise contrasts (FDR-corrected).

### Linguistic Focus Test

The linguistic focus test (schematic Fig. 9A) measured participants’ ability to detect where a contrastive accent was placed in a sentence, based on only one type of acoustic dimension (F0 or Duration) or both combined (as in natural speech). As shown in Fig. 10B and Table 2, overall both groups performed best when they heard F0 and duration cues together, worst when only duration cues were present, and in between when there were only F0 cues (main effect of Condition χ^2^(4) = 168.4, p < 0.001). This suggests that both groups benefitted from redundant information, and that F0 was more useful for detecting focus than duration, in line with results from Experiment 1. On the whole, controls performed more accurately than amusics (main effect of Group χ^2^(3) = 14.63, *p* = 0.002). However, the two groups were differentially affected by whether F0 or duration cues were present in the stimuli (interaction of Group X Condition χ^2^(2) = 12.05, p = 0.002). When relying on duration alone, amusics performed similarly to controls, but when they needed to rely on F0 they performed significantly less accurately (*p*=.019; Table 2). This disadvantage held where F0 was the sole cue, as well as in the Combined F0+duration cue condition. This result stands in contrast to that in the musical phrase test, where performance in the combined (pitch+duration) condition did not differ between groups.

**Table 2:** Linguistic Focus test: pairwise comparisons of marginal means (p-values FDR adjusted).

| **Condition** | **Group** | **Contrast** | **OR** | **SE** | **Z** | **p** |
| --- | --- | --- | --- | --- | --- | --- |
| Combined | ~ | CONT vs AMUS | 2.44 | 0.13 | **2.71** | **0.009** |
| Duration | ~ | CONT vs AMUS | 1.11 | 0.24 | 0.39 | 0.697 |
| F0 | ~ | CONT vs AMUS | 2.00 | 0.14 | **2.39** | **0.019** |
| ~ | AMUS | Combined vs F0 | 2.06 | 0.37 | **4.01** | **<.001** |
| ~ | AMUS | Combined vs Duration | 3.71 | 0.64 | **7.56** | **<.001** |
| ~ | AMUS | F0 vs Duration | 1.80 | 0.28 | **3.83** | **<.001** |
| ~ | CONT | Combined vs F0 | 2.52 | 0.62 | **3.77** | **<.001** |
| ~ | CONT | Combined vs Duration | 8.15 | 1.84 | **9.31** | **<.001** |
| ~ | CONT | F0 vs Duration | 3.23 | 0.57 | **6.65** | **<.001** |
| ~ | ALL | Combined vs F0 | 2.28 | 0.36 | **5.22** | **<.001** |
| ~ | ALL | Combined vs Duration | 5.50 | 0.82 | **11.44** | **<.001** |
| ~ | ALL | F0 vs Duration | 2.41 | 0.30 | **7.10** | **<.001** |

## Linguistic Phrase Test

The Linguistic Phrase Perception Test (schematic Fig 9B) measured participants’ ability to detect phrase boundaries in speech which are cued by F0 only, duration only, or both F0 and duration. Cue type affected performance across groups (main effect of Condition χ^2^(4) = 83.06, p < 0.001). Participants performed least accurately when they had to rely on F0 cues alone, better when they relied on duration alone, and most accurately when both F0 and duration were present together (see Fig. 10B and Table 3). This is consistent with prior evidence suggesting that duration is a more reliable cue for the detection of phrase boundaries than pitch (Streeter, 1978). As in the Focus test, redundant cues benefitted both groups, but in contrast to the pattern in the Focus Test, duration was a more reliable cue to linguistic phrase boundary perception than pitch.

**Table 3:** Post hoc contrasts, Linguistic Phrase Test.

| **Condition** | **Group** | **Contrast** | **OR** | **SE** | **Z** | **P** |
| --- | --- | --- | --- | --- | --- | --- |
| Combined | ~ | CONT vs AMUS | 1.10 | 0.26 | 0.32 | 0.841 |
| Duration | ~ | CONT vs AMUS | 1.01 | 0.27 | 0.02 | 0.985 |
| F0 | ~ | CONT vs AMUS | 1.35 | 0.20 | 1.12 | 0.338 |
| ~ | AMUS | Combined vs F0 | 2.88 | 0.44 | **7.00** | **<0.001** |
| ~ | AMUS | Combined vs Duration | 1.77 | 0.28 | **3.64** | **0.001** |
| ~ | AMUS | Duration vs Pitch | 1.63 | 0.08 | **3.56** | **0.001** |
| ~ | CONT | Combined vs F0 | 2.34 | 0.37 | **5.37** | **<0.001** |
| ~ | CONT | Combined vs Duration | 1.93 | 0.31 | **4.10** | **<0.001** |
| ~ | CONT | Duration vs F0 | 1.21 | 0.12 | 1.34 | 0.268 |
| ~ | ALL | Combined vs F0 | 2.60 | 0.28 | **8.71** | **<.001** |
| ~ | ALL | Combined vs Duration | 1.85 | 0.21 | **5.48** | **<.001** |
| ~ | ALL | Duration vs F0 | 1.41 | 0.07 | **3.43** | **<.001** |

Amusics did not differ significantly from controls in overall accuracy (main effect of Group χ^2^(3) = 2.69, *p* **=** 0.44) nor was the groups’ performance significantly differently affected by which acoustic cues were present (interaction of Group X Condition χ^2^(2) = 2.33, p = 0.31). Because we had hypothesized *a priori* that amusics would rely more on duration than F0 (and that controls would show similar performance across the two conditions), we conducted pairwise contrasts to test this prediction. Amusics did indeed show significantly greater accuracy with duration than with F0 cues (*p*=.001; Table 3), whereas controls did not. (For completeness, all other (post-hoc) pairwise comparisons are also reported).

## Discussion

Music and speech carry information across multiple acoustic dimensions, with distinct dimensions often providing redundant information to support perceptual decisions. We tested whether this multidimensional redundancy makes music and speech robust to individual differences in perceptual abilities. We found that amusics perceived both speech and music well when they could rely on a redundant, unimpaired channel (duration), either on its own, or together with F0 or pitch. Furthermore, for musical and linguistic phrase perception, amusics were able to achieve equivalent performance to control participants when boundaries were redundantly cued by F0/pitch and duration. Thus, redundancy in the production of communicative signals makes possible robust message transmission in the face of individual differences in auditory perception.

Musical aptitude is often measured with tests that target specific domains like perception of melody or rhythm (Gordon, 2002; Wallentin, Nielsen, Friis-Olivarius, Vuust, & Vuust, 2010)). This is, however, unlike actual music listening in the real world. Naturally produced musical structures, such as musical phrases, are often conveyed by simultaneous (i.e. redundant) cues (Palmer & Krumhansl, 1987)). Here, we find no evidence that amusics’ intuitions about musical phrase structure are impaired as long as durational cues are present. Previous studies of musical phrase judgments have found that duration and pitch cues carry equal weights, without additional benefits from being able to combine the two (Palmer & Krumhansl, 1987)), a finding we replicated in our control participants. Amusics, on the other hand, showed a gain from redundancy. While amusics may not fully appreciate aspects of music that relate to pitch, we show that they can parse musical structures when another relevant cue is available. Future work should investigate whether this spared musical perception in amusics extends to other musical features which are communicated by redundant cues, such as musical beat perception (Hannon, Snyder, Eerola, & Krumhansl, 2004)).

One possible outcome was that amusics would show superior duration processing that they had developed to compensate for their pitch deficit. The data here do not support this. The amusics showed similar - and not significantly more accurate - duration perception ability compared to controls across the music and language tests, as well as similar psychophysical duration discrimination thresholds. The present data suggest that rather than developing exceptional duration processing ability, all that may be necessary is a re-weighting in perception to emphasize dimensions for which perception is more accurate.

## General Discussion

There is a remarkable diversity in the use and weighting of perceptual dimensions in speech and music in the typical population (Chandrasekaran et al., 2010; Hazan & Rosen, 1991; Idemaru et al., 2012; Kim et al., 2018; Kong & Edwards, 2016; Schertz, Cho, Lotto, & Warner, 2015b). How do such stable individual differences arise, and how do communication systems facilitate robust information transmission despite this variability? We hypothesized that the relative facility that an individual listener shows in extracting information from perceptual dimensions would bias the relative weighting of these dimensions when they are used to convey communicative content. We tested this with an ‘experiment of nature’ - people with congenital difficulties in processing pitch, but not duration - and asked whether such a specific perceptual starting state might bias these listeners away from using their unreliable dimension (pitch) and toward another that is processed more reliably (duration cues) during perceptual categorization. We also asked whether such biases would be specific to cases in which the unreliable dimension (pitch) tends to be the primary cue - for instance in prosodic emphasis - or if it would extend to cases in which it was a secondary cue (e.g., categorization of onset syllable voicing). Finally, we examined whether the existence of redundant cues in speech and music (i.e., the presence of both durational and pitch cues) makes possible relatively robust transmission of message-level information, despite profound individual differences in perceptual acuity.

We first asked the question of whether one’s relative ability to perceive certain auditory dimensions is indeed linked to perceptual weightings of those dimensions in speech perception. In the case of prosodic emphasis, typical listeners tend to rely mainly on F0, but people with pitch perception difficulties tended to rely less on F0. When F0 and duration cues conflicted, amusic participants generally categorized stimuli according to duration cues, whereas the control group tended to categorize the very same stimuli according to F0 cues. For onset syllable voicing (phonetic categorization, for which F0 is typically a secondary dimension), on the other hand, there were no significant differences between people with pitch perception difficulties and control participants. Importantly, the F0 frequency range used to convey information across both of these tasks was considerably greater than the pitch discrimination thresholds of each participant. Thus, although the participants with pitch perception difficulties could have used pitch perception for prosodic categorization, they may have learned over time that pitch cues conveyed by F0 contours were less useful for them compared to temporal/durational cues. Alternatively, such a reweighting away from use of F0 cues in this case may reflect a strategic (but unconscious) use of information that is better perceived. An analogy here is that of a television viewer who can watch equivalent six o’clock news programs on Channel 1 or Channel 2. Channel 2 can be received, but only with poor quality. However, they have perfect reception for Channel 1, and therefore develop a bias to watch the news there.

Next, we asked whether the redundant information conveyed by pitch/F0 and duration is sufficient to enable reliable message transmission via prosody despite severe perceptual deficits. In a follow-up experiment, we found that, although participants with pitch perception difficulties struggled to decode musical and prosodic information when conveyed only via pitch and F0 cues, they performed equivalently to control participants when durational cues were added. Thus, the existence of redundant cues to structure in speech and music ensures that these systems are robust even to extreme perceptual deficits. Because of the redundancy built into speech and music, people with specific and even severe auditory deficits can perceive structure nonetheless.

To keep the experimental design simple, we only examined two auditory dimensions -- pitch changes conveyed through F0, where we suspected our groups would show a difference, and duration, where we believed they would not. Outside the laboratory there are other cues that individuals could take advantage of, such as vowel quality, which is also associated with phrase boundaries, and pitch accents (Sluijter & van Heuven, 1996; Streeter, 1978). Accents also carry visual correlates, such as head movements, beat gestures, and eyebrow raises (e.g. Beskow, Granström, Conference, 2006, 2006; Flecha-García, 2010; Krahmer & Swerts, 2007), which individuals may also be able to use to compensate for their pitch impairment in audiovisual speech perception. Moreover, top-down processes such as the use of lexical knowledge can also help disambiguate unclear speech (Connine & Clifton, 1987; Ganong, 1980) and talker identity cues from the visual modality help listeners to disambiguate acoustic-phonetic cues (Zhang & Holt, 2018). Individuals may be able to modify the extent to which they make use of any or all of these different sources of information in response to their idiosyncratic set of strengths and weaknesses. For example, individuals with widespread auditory processing problems may rely more heavily on top-down lexical information, or visual cues.

Our participants were all adults, and so could take advantage of decades of experience in perceiving speech. One open question is the length of time necessary for developing perceptual categorization strategies which take into account the reliability of perception. For example, although correlations between auditory perception thresholds and speech perception are generally weak or nonexistent in adults listening to their native language, stronger correlations have been reported in young children (Bavin, Grayden, Scott, & Stefanakis, 2010; Boets, Wouters, van Wieringen, De Smedt, & Ghesquière, 2008) and adults in the initial stages of learning a foreign language (Chandrasekaran, Kraus, & Wong, 2012). Auditory processing may, therefore, be a bottleneck for speech perception in the initial stages of acquiring a language, at which time listeners have not yet developed the ability to flexibly switch between cues, or have developed stable adult-like perceptual weights across acoustic dimensions (Idemaru & Holt, 2013; Mayo & Turk, 2004). In later stages of language acquisition, correlations between auditory processing and speech perception may dissipate, although language delays due to earlier problems with speech may persist Rosen:2003cy, Ramus:2013ja}. This hypothesis could be tested by examining the relationship between cue weighting and auditory processing in children. For instance, children might show a smaller or nonexistent tendency to rely more on the dimensions which they are better able to process.

That individuals can compensate for perceptual impairments through greater reliance on higher-fidelity auditory dimensions, along with evidence that perceptual weighting strategies can be altered through training (Francis et al., 2008; Lim & Holt, 2011; Wu et al., 2016, suggests that it may be possible to ameliorate speech perception deficits via the development of targeted auditory processing training batteries. In particular, participants could be trained to rely more heavily on the attributes which they can more easily track. For example, participants with temporal perception deficits could be trained to focus on pitch-based cues to syntactic and phonological structure.

The extent to which individual differences in dimensional weighting are stable across tasks and individuals remains an open question. As discussed above, amusics rely less on F0 in prosodic but not phonetic categorization, suggesting that in some cases dimensional weighting can vary. Future work could examine whether amusics also rely less on pitch information when detecting musical structure. The boundaries of musical phrases, for example, are conveyed by pitch and durational changes similar to those found at the boundaries of linguistic phrases, including phrase-final lengthening and changes in pitch (Tierney, Russo, & Patel, 2011). Accordingly, typical listeners listen for both pitch and durational cues when judging the completeness of a musical phrase (Jusczyk & Krumhansl, 1993; Palmer & Krumhansl, 1987). Similarly, the location of musical beats can be conveyed by lengthening of the note on which the beat falls and/or a change in melodic contour (Ellis & Jones, 2009; Hannon et al., 2004; Prince, 2014). Despite the existence of multiple redundant cues for musical structure, dimensional weighting in music perception has not been systematically investigated, and it remains unknown whether individual differences in musical dimensional weighting exist, and whether listeners will dynamically alter their musical dimensional weighting in response to short-term changes in cue distribution.

Our finding that amusics can minimize the impact of their perceptual impairment by focusing on preserved cues suggests that other populations may be able to take advantage of a similar strategy. Individuals with autism (O’Connor, 2012), ADHD (Riccio, Hynd, Cohen, Hall, & Molt, 1994), and beat deafness (Phillips-Silver et al., 2011), for example, have been reported to display impaired temporal but not pitch perception. Our model population was able to integrate pitch and duration together to perform the tasks, and this strategy may limit the impact of impaired auditory perception in some of these populations. Other groups, however, might find this strategy less successful; for instance individuals with autism have difficulty integrating information across multiple senses (Marco, Hinkley, Hill, & Nagarajan, 2011).

In conclusion, our results showcase how widespread competency can hide individual differences in how individuals perceive the world. Perception can, on the surface, appear to be seamless and universal, with most people appearing to arrive at the same interpretations from the same information. This, however, can mask the true diversity of human experience.

## Contributions

A.T.T. developed the study concept. All authors contributed to the design. K.J. performed testing, data collection, data analysis and drafted the manuscript. F.D., A.T.T. and L.H. provided critical revisions. All authors approved the final version of the manuscript for submission.

## Acknowledgments

We thank Stuart Rosen, Marcus Pearce, Laura Staum-Casasanto, Alex Martin, Aniruddh Patel, Clare Press and Lauren Stewart for helpful comments and discussion. We also thank all our participants. The work was funded by a Wellcome Trust Seed Award #109719/Z/15/Z to A.T.T., a Reg and Molly Buck Award from SEMPRE to K.J., and a Leverhulme Trust Early Career Fellowship to K.J.

## Data Availability

The data that support the findings of this study are available from the corresponding author upon reasonable request.

## References

Anwyl-Irvine, A., Massonnié, J., Flitton, A., Kirkham, N., & Evershed, J. (2018). Gorillas in our Midst: Gorilla.sc, a new web-based Experiment Builder, 1–19. http://doi.org/10.1101/438242

Ayotte, J., Peretz, I., & Hyde, K. (2002). Congenital amusiaA group study of adults afflicted with a music-specific disorder. Brain, 125(2), 238–251. http://doi.org/10.1093/brain/awf028

Bavin, E. L., Grayden, D. B., Scott, K., & Stefanakis, T. (2010). Testing Auditory Processing Skills and their Associations with Language in 4—5-year-olds. Language and Speech, 53(1), 31–47. http://doi.org/10.1177/0023830909349151

Bench, J., Kowal, Å., & Bamford, J. (1979). The Bkb (Bamford-Kowal-Bench) Sentence Lists for Partially-Hearing Children, 13(3), 108–112. http://doi.org/10.3109/03005367909078884

Bentrovato, S., Devescovi, A., D’Amico, S., Wicha, N., & Bates, E. (2003). The Effect of Grammatical Gender and Semantic Context on Lexical Access in Italian Using a Timed Word-Naming Paradigm. Journal of Psycholinguistic Research, 32(4), 417–430. http://doi.org/10.1023/A:1024899513147

Beskow, J., Granström, B., Conference, D. H. N. I., 2006. (2006). Visual correlates to prominence in several expressive modes. Ninth International Conference on Spoken Language Processing.

Boebinger, D., Evans, S., Rosen, S., Lima, C. F., Manly, T., & Scott, S. K. (2015). Musicians and non-musicians are equally adept at perceiving masked speech. The Journal of the Acoustical Society of America, 137(1), 378–387. http://doi.org/10.1121/1.4904537

Boersma, P. (2002). Praat: a system for doing phonetics by computer. Glot International, 5.

Boets, B., Wouters, J., van Wieringen, A., De Smedt, B., & Ghesquière, P. (2008). Modelling relations between sensory processing, speech perception, orthographic and phonological ability, and literacy achievement. Brain and Language, 106(1), 29–40. http://doi.org/10.1016/j.bandl.2007.12.004

Breen, M., Fedorenko, E., Wagner, M., & Gibson, E. (2010). Acoustic correlates of information structure. Language and Cognitive Processes, 25(7-9), 1044–1098. http://doi.org/10.1080/01690965.2010.504378

Chandrasekaran, B., Kraus, N., & Wong, P. C. M. (2012). Human inferior colliculus activity relates to individual differences in spoken language learning. J Neurophysiol. http://doi.org/10.1152/jn.00923.2011;page:string:Article/Chapter

Chandrasekaran, B., Sampath, P. D., & Wong, P. C. M. (2010). Individual variability in cue-weighting and lexical tone learning. The Journal of the Acoustical Society of America, 128(1), 456–465. http://doi.org/10.1121/1.3445785

Chrabaszcz, A., Winn, M., Lin, C. Y., & Idsardi, W. J. (2014). Acoustic Cues to Perception of Word Stress by English, Mandarin, and Russian Speakers. Journal of Speech, Language, and Hearing Research, 57(4), 1468–1479. http://doi.org/10.1044/2014_JSLHR-L-13-0279

Connine, C. M., & Clifton, C. (1987). Interactive use of lexical information in speech perception. Journal of Experimental Psychology: Human Perception and Performance, 13(2), 291–299. http://doi.org/10.1037//0096-1523.13.2.291

Dalla Bella, S., Giguère, J.-F., & Peretz, I. (2009). Singing in congenital amusia. The Journal of the Acoustical Society of America, 126(1), 414–424. http://doi.org/10.1121/1.3132504

de Pijper, J. R., & Sanderman, A. A. (1994). On the perceptual strength of prosodic boundaries and its relation to suprasegmental cues. The Journal of the Acoustical Society of America, 96(4), 2037–2047. http://doi.org/10.1121/1.410145

Ellis, R. J., & Jones, M. R. (2009). The role of accent salience and joint accent structure in meter perception. Journal of Experimental Psychology: Human Perception and Performance, 35(1), 264–280. http://doi.org/10.1037/a0013482

Elman, J. L. (2009). On the Meaning of Words and Dinosaur Bones: Lexical Knowledge Without a Lexicon. Cognitive Science, 33(4), 547–582. http://doi.org/10.1111/j.1551-6709.2009.01023.x

Fear, B. D., Cutler, A., & Butterfield, S. (1995). The strong/weak syllable distinction in English. The Journal of the Acoustical Society of America, 97(3), 1893–1904. http://doi.org/10.1121/1.412063

Flecha-García, M. L. (2010). Eyebrow raises in dialogue and their relation to discourse structure, utterance function and pitch accents in English. Speech Communication, 52(6), 542–554. http://doi.org/10.1016/j.specom.2009.12.003

Francis, A. L., Baldwin, K., & Nusbaum, H. C. (2000). Effects of training on attention to acoustic cues. Perception & Psychophysics, 62(8), 1668–1680. http://doi.org/10.3758/BF03212164

Francis, A. L., Kaganovich, N., & Driscoll-Huber, C. (2008). Cue-specific effects of categorization training on the relative weighting of acoustic cues to consonant voicing in English. The Journal of the Acoustical Society of America, 124(2), 1234–1251. http://doi.org/10.1121/1.2945161

Ganong, W. F. (1980). Phonetic categorization in auditory word perception. Journal of Experimental Psychology: Human Perception and Performance, 6(1), 110–125. http://doi.org/10.1037//0096-1523.6.1.110

Gordon, E. E. (2002). Primary Measures of Music Audiation.

Grassi, M., & Soranzo, A. (2009). MLP: A MATLAB toolbox for rapid and reliable auditory threshold estimation. Behavior Research Methods, 41(1), 20–28. http://doi.org/10.3758/BRM.41.1.20

Haggard, M., Ambler, S., & Callow, M. (1970). Pitch as a Voicing Cue. The Journal of the Acoustical Society of America, 47(2B), 613–617. http://doi.org/10.1121/1.1911936

Hannon, E. E., Snyder, J. S., Eerola, T., & Krumhansl, C. L. (2004). The Role of Melodic and Temporal Cues in Perceiving Musical Meter. Journal of Experimental Psychology: Human Perception and Performance, 30(5), 956–974. http://doi.org/10.1037/0096-1523.30.5.956

Hazan, V., & Rosen, S. (1991). Individual variability in the perception of cues to place contrasts in initial stops. Perception & Psychophysics, 49(2), 187–200. http://doi.org/10.3758/BF03205038

Holt, L. L., & Lotto, A. J. (2006). Cue weighting in auditory categorization: Implications for first and second language acquisition. The Journal of the Acoustical Society of America, 119(5), 3059–3071. http://doi.org/10.1121/1.2188377

Holt, L. L., & Lotto, A. J. (2008). Speech Perception Within an Auditory Cognitive Science Framework. Current Directions in Psychological Science, 17(1), 42–46. http://doi.org/10.1111/j.1467-8721.2008.00545.x

Holt, L. L., Tierney, A. T., Guerra, G., Laffere, A., & Dick, F. (2018). Dimension-selective attention as a possible driver of dynamic, context-dependent re-weighting in speech processing. Hearing Research, 366, 50–64. http://doi.org/10.1016/j.heares.2018.06.014

Hutchins, S., Gosselin, N., & Peretz, I. (2010). Identification of Changes along a Continuum of Speech Intonation is Impaired in Congenital Amusia. Frontiers in Psychology, 1. http://doi.org/10.3389/fpsyg.2010.00236

Hyde, K. L., & Peretz, I. (2004). Brains That Are out of Tune but in Time. Psychological Science, 15(5), 356–360. http://doi.org/10.1111/j.0956-7976.2004.00683.x

Idemaru, K., & Holt, L. L. (2011). Word recognition reflects dimension-based statistical learning. Journal of Experimental Psychology: Human Perception and Performance, 37(6), 1939–1956. http://doi.org/10.1037/a0025641

Idemaru, K., & Holt, L. L. (2014). Specificity of dimension-based statistical learning in word recognition. Journal of Experimental Psychology: Human Perception and Performance, 40(3), 1009–1021. http://doi.org/10.1037/a0035269

Idemaru, K., Holt, L. L., & Seltman, H. (2012). Individual differences in cue weights are stable across time: The case of Japanese stop lengths. The Journal of the Acoustical Society of America, 132(6), 3950–3964. http://doi.org/10.1121/1.4765076

Jiang, C., Hamm, J. P., Lim, V. K., Kirk, I. J., & Yang, Y. (2010). Processing melodic contour and speech intonation in congenital amusics with Mandarin Chinese. Neuropsychologia, 48(9), 2630–2639. http://doi.org/10.1016/j.neuropsychologia.2010.05.009

Jusczyk, P. W., & Krumhansl, C. L. (1993). Pitch and rhythmic patterns affecting infants’ sensitivity to musical phrase structure. Journal of Experimental Psychology: Human Perception and Performance, 19(3), 627–640. http://doi.org/10.1037/0096-1523.19.3.627

Karlin, J. E. (1942). A factorial study of auditory function. Psychometrika, 7(4), 251–279. http://doi.org/10.1007/BF02288628

Kawahara, H., & Irino, T. (2005). Underlying Principles of a High-quality Speech Manipulation System STRAIGHT and Its Application to Speech Segregation.In Speech Separation by Humans and Machines (pp. 167–180). Boston: Kluwer Academic Publishers. http://doi.org/10.1007/0-387-22794-6_11

Kidd, G. R., Watson, C. S., & Gygi, B. (2007). Individual differences in auditory abilities. The Journal of the Acoustical Society of America, 122(1), 418–435. http://doi.org/10.1121/1.2743154

Kim, D., Clayards, M., & Goad, H. (2018). A longitudinal study of individual differences in the acquisition of new vowel contrasts. Journal of Phonetics, 67, 1–20. http://doi.org/10.1016/j.wocn.2017.11.003

Kjelgaard, M. M., & Speer, S. R. (1999). Prosodic Facilitation and Interference in the Resolution of Temporary Syntactic Closure Ambiguity. Journal of Memory and Language, 40(2), 153–194. http://doi.org/10.1006/jmla.1998.2620

Kong, E. J., & Edwards, J. (2016). Individual differences in categorical perception of speech: Cue weighting and executive function. Journal of Phonetics, 59, 40–57. http://doi.org/10.1016/j.wocn.2016.08.006

Krahmer, E., & Swerts, M. (2007). The effects of visual beats on prosodic prominence: Acoustic analyses, auditory perception and visual perception. Journal of Memory and Language, 57(3), 396–414. http://doi.org/10.1016/j.jml.2007.06.005

Lerdahl, F., & Jackendoff, R. (1985). A Generative Theory of Tonal Music. MIT Press.

Liberman, A. M., & Mattingly, I. G. (1985). The motor theory of speech perception revised. Cognition, 21(1), 1–36. http://doi.org/10.1016/0010-0277(85)90021-6

Liberman, A. M., Cooper, F. S., Shankweiler, D. P., & Studdert-Kennedy, M. (1967). Perception of the Speech Code. Psychological Review, 74(6), 431–461.

Lim, S.-J., & Holt, L. L. (2011). Learning Foreign Sounds in an Alien World: Videogame Training Improves Non-Native Speech Categorization. Cognitive Science, 35(7), 1390–1405. http://doi.org/10.1111/j.1551-6709.2011.01192.x

Lisker, L. (1957). Closure Duration and the Intervocalic Voiced-Voiceless Distinction in English. Language, 33(1), 42. http://doi.org/10.2307/410949

Lisker, L. (2016). “Voicing” in English: A Catalogue of Acoustic Features Signaling /b/ Versus /p/ in Trochees. Language and Speech, 29(1), 3–11. http://doi.org/10.1177/002383098602900102

Liu, F., Jiang, C., Wang, B., Xu, Y., & Patel, A. D. (2015). A music perception disorder (congenital amusia) influences speech comprehension. Neuropsychologia, 66(C), 111–118. http://doi.org/10.1016/j.neuropsychologia.2014.11.001

Liu, F., Patel, A. D., Fourcin, A., & Stewart, L. (2010). Intonation processing in congenital amusia: discrimination, identification and imitation. Brain, 133(6), 1682–1693. http://doi.org/10.1093/brain/awq089

Liu, R., & Holt, L. L. (2015). Dimension-based statistical learning of vowels. Journal of Experimental Psychology: Human Perception and Performance, 41(6), 1783–1798. http://doi.org/10.1037/xhp0000092

Lu, Y., & Cooke, M. (2009). The contribution of changes in F0 and spectral tilt to increased intelligibility of speech produced in noise. Speech Communication, 51(12), 1253–1262. http://doi.org/10.1016/j.specom.2009.07.002

Lutfi, R. A., & Liu, C.-J. (2007). Individual differences in source identification from synthesized impact sounds. The Journal of the Acoustical Society of America, 122(2), 1017–1028. http://doi.org/10.1121/1.2751269

Marco, E. J., Hinkley, L. B. N., Hill, S. S., & Nagarajan, S. S. (2011). Sensory Processing in Autism: A Review of Neurophysiologic Findings. Pediatric Research, 69(5 Part 2), 48R–54R. http://doi.org/10.1203/PDR.0b013e3182130c54

Massaro, D. W., & Cohen, M. M. (1976). The contribution of fundamental frequency and voice onset time to the /zi/-/si/ distinction. The Journal of the Acoustical Society of America, 60(3), 704–717. http://doi.org/10.1121/1.381143

Massaro, D. W., & Cohen, M. M. (1977). Voice onset time and fundamental frequency as cues to the /zi/-/si/ distinction. Perception & Psychophysics, 22(4), 373–382. http://doi.org/10.3758/BF03199703

Mattys, S. L. (2000). The perception of primary and secondary stress in English. Perception & Psychophysics, 62(2), 253–265. http://doi.org/10.3758/BF03205547

McMurray, B., & Jongman, A. (2011). What information is necessary for speech categorization? Harnessing variability in the speech signal by integrating cues computed relative to expectations. Psychological Review, 118(2), 219–246. http://doi.org/10.1037/a0022325

McMurray, B., Aslin, R. N., Tanenhaus, M. K., Spivey, M. J., & Subik, D. (2008). Gradient sensitivity to within-category variation in words and syllables. Journal of Experimental Psychology: Human Perception and Performance, 34(6), 1609–1631. http://doi.org/10.1037/a0011747

Mignault Goulet, G., Moreau, P., Robitaille, N., & Peretz, I. (2012). Congenital Amusia Persists in the Developing Brain after Daily Music Listening. PLoS ONE, 7(5), e36860. http://doi.org/10.1371/journal.pone.0036860

Moreau, P., Jolicœur, P., & Peretz, I. (2013). Pitch discrimination without awareness in congenital amusia: Evidence from event-related potentials. Brain and Cognition, 81(3), 337–344. http://doi.org/10.1016/j.bandc.2013.01.004

Nan, Y., Sun, Y., & Peretz, I. (2010). Congenital amusia in speakers of a tone language: association with lexical tone agnosia. Brain, 133(9), 2635–2642. http://doi.org/10.1093/brain/awq178

O’Connor, K. (2012). Auditory processing in autism spectrum disorder: A review. Neuroscience and Biobehavioral Reviews, 36(2), 836–854. http://doi.org/10.1016/j.neubiorev.2011.11.008

Palmer, C., & Krumhansl, C. L. (1987). Independent temporal and pitch structures in determination of musical phrases. Journal of Experimental Psychology: Human Perception and Performance, 13(1), 116–126. http://doi.org/10.1037//0096-1523.13.1.116

Patel, A. D. (2014). Can nonlinguistic musical training change the way the brain processes speech? The expanded OPERA hypothesis. Hearing Research, 308, 98–108. http://doi.org/10.1016/j.heares.2013.08.011

Patel, A. D., Foxton, J. M., & Griffiths, T. D. (2005). Musically tone-deaf individuals have difficulty discriminating intonation contours extracted from speech. Brain and Cognition, 59(3), 310–313. http://doi.org/10.1016/j.bandc.2004.10.003

Patel, A. D., Wong, M., Foxton, J., Lochy, A., & Peretz, I. (2008). Speech intonation perceptuion deficits in musical tone deafness (congenital amusia). Music Perception: an Interdisciplinary Journal, 25(4), 357–368. http://doi.org/10.1525/mp.2008.25.4.357

Peretz, I., & Vuvan, D. T. (2017). Prevalence of congenital amusia. European Journal of Human Genetics, 25(5), 625–630. http://doi.org/10.1038/ejhg.2017.15

Peretz, I., Ayotte, J., Zatorre, R. J., Mehler, J., Neuron, P. A., Penhune, V. B., & Jutras, B. (2002). Congenital amusia: a disorder of fine-grained pitch discrimination. Neuron, 33(2), 185–191. http://doi.org/10.1016/S0896-6273(01)00580-3

Peretz, I., Brattico, E., & Tervaniemi, M. (2005). Abnormal electrical brain responses to pitch in congenital amusia. Annals of Neurology, 58(3), 478–482. http://doi.org/10.1002/ana.20606

Peretz, I., Champod, A. S., & Hyde, K. (2003). Varieties of Musical Disorders. Annals of the New York Academy of Sciences, 999(1), 58–75. http://doi.org/10.1196/annals.1284.006

Phillips-Silver, J., Toiviainen, P., Gosselin, N., Piché, O., Nozaradan, S., Palmer, C., & Peretz, I. (2011). Born to dance but beat deaf: A new form of congenital amusia. Neuropsychologia, 49(5), 961–969. http://doi.org/10.1016/j.neuropsychologia.2011.02.002

Prince, J. B. (2014). Contributions of pitch contour, tonality, rhythm, and meter to melodic similarity. Journal of Experimental Psychology: Human Perception and Performance, 40(6), 2319–2337. http://doi.org/10.1037/a0038010

Riccio, C. A., Hynd, G. W., Cohen, M. J., Hall, J., & Molt, L. (1994). Comorbidity of Central Auditory Processing Disorder and Attention-Deficit Hyperactivity Disorder. Journal of the American Academy of Child & Adolescent Psychiatry, 33(6), 849–857. http://doi.org/10.1097/00004583-199407000-00011

Schaffrath, H., & Park, D. H. M. (1995). The Essen folksong collection in kern format.[computer database].

Schertz, J., Cho, T., Lotto, A., & Warner, N. (2015a). Individual differences in perceptual adaptability of foreign sound categories. Attention, Perception, & Psychophysics, 78(1), 355–367. http://doi.org/10.3758/s13414-015-0987-1

Schertz, J., Cho, T., Lotto, A., & Warner, N. (2015b). Individual differences in phonetic cue use in production and perception of a non-native sound contrast. Journal of Phonetics, 52, 183–204. http://doi.org/10.1016/j.wocn.2015.07.003

Sluijter, A. M. C., & van Heuven, V. J. (1996). Acoustic correlates of linguistic stress and accent in Dutch and American English (Vol. 2, pp. 630–633). Presented at the Proceeding of Fourth International Conference on Spoken Language Processing. ICSLP’96, IEEE. http://doi.org/10.1109/ICSLP.1996.607440

Stankov, L., & Horn, J. L. (1980). Human abilities revealed through auditory tests. Journal of Educational Psychology, 72(1), 21–44. http://doi.org/10.1037/0022-0663.72.1.21

Streeter, L. A. (1978). Acoustic determinants of phrase boundary perception. The Journal of the Acoustical Society of America, 64(6), 1582–1592. http://doi.org/10.1121/1.382142

Surprenant, A. M., & Watson, C. S. (2001). Individual differences in the processing of speech and nonspeech sounds by normal-hearing listeners. The Journal of the Acoustical Society of America, 110(4), 2085–2095. http://doi.org/10.1121/1.1404973

Tierney, A. T., Russo, F. A., & Patel, A. D. (2011). The motor origins of human and avian song structure. Proceedings of the National Academy of Sciences, 108(37), 15510–15515. http://doi.org/10.1073/pnas.1103882108

Toscano, J. C., & McMurray, B. (2010). Cue Integration With Categories: Weighting Acoustic Cues in Speech Using Unsupervised Learning and Distributional Statistics. Cognitive Science, 34(3), 434–464. http://doi.org/10.1111/j.1551-6709.2009.01077.x

Utman, J. A., Blumstein, S. E., & Burton, M. W. (2000). Effects of subphonetic and syllable structure variation on word recognition. Perception & Psychophysics, 62(6), 1297–1311. http://doi.org/10.3758/BF03212131

Vuvan, D. T., Nunes-Silva, M., & Peretz, I. (2015). Meta-analytic evidence for the non-modularity of pitch processing in congenital amusia. Cortex, 69(C), 186–200. http://doi.org/10.1016/j.cortex.2015.05.002

Wallentin, M., Nielsen, A. H., Friis-Olivarius, M., Vuust, C., & Vuust, P. (2010). The Musical Ear Test, a new reliable test for measuring musical competence. Learning and Individual Differences, 20(3), 188–196. http://doi.org/10.1016/j.lindif.2010.02.004

Watson, C. S., & Kidd, G. R. (2002). On the Lack of Association between Basic Auditory Abilities, Speech Processing, and other Cognitive Skills. Seminars in Hearing, 23(1), 083–094. http://doi.org/10.1055/s-2002-24978

Watson, C. S., Jensen, J. K., Foyle, D. C., Leek, M. R., & Goldgar, D. E. (1982). Performance of 146 normal adult listeners on a battery of auditory discrimination tests. The Journal of the Acoustical Society of America, 71(S1), S73–S73. http://doi.org/10.1121/1.2019533

Winn, M. B., Chatterjee, M., & Idsardi, W. J. (2013). Roles of Voice Onset Time and F0 in Stop Consonant Voicing Perception: Effects of Masking Noise and Low-Pass Filtering. Journal of Speech, Language, and Hearing Research, 56(4), 1097–1107. http://doi.org/10.1044/1092-4388(2012/12-0086)

Winter, B. (2014). Spoken language achieves robustness and evolvability by exploiting degeneracy and neutrality. BioEssays, 36(10), 960–967. http://doi.org/10.1002/bies.201400028

Wu, Y. C., & Holt, L. L. (2018). Phonetic category activation drives dimension-based adaptive tuning in speech perception. Proceedings of the Cognitive Science Society.

Yu, A., & Zellou, G. (2019). Individual Differences in Language Processing: Phonology. Anual Review of Linguistics, 5, 6.1–6.20. http://doi.org/10.1146/annurev-linguistics-011516-033815

